# Systemic Cysteine Elevation Sustains T-Cell Activation to Potentiate PD-1 Blockade

**DOI:** 10.64898/2026.03.16.711688

**Authors:** Xiaoshuang Wang, Zi Wang, Yuqi Guo, Fangxi Xu, Jinhui Zhu, Scott C. Thomas, Deepak Saxena, Jinghang Xie, Xin Li

**Affiliations:** Plastic Surgery, Maxillofacial & Oral Health Department, University of Virginia School of Medicine, Charlottesville, VA 22903, USA; Molecular Pathobiology Department, New York University College of Dentistry, New York, NY 10010, USA; Molecular Physiology and Biological Physics Department, University of Virginia School of Medicine, Charlottesville, VA 22903, USA; Laura and Isaac Perlmutter Cancer Center, NYU Grossman School of Medicine, New York, NY 10016, USA; Department of Surgery, NYU Grossman School of Medicine, New York, NY 10016, USA; UVA Comprehensive Cancer Center, University of Virginia School of Medicine, Charlottesville, VA 22903, USA; Neuroscience Department, University of Virginia School of Medicine, Charlottesville, VA 22903, USA

**Keywords:** Probiotics, anti-PD-1, T cell, Cystine, Cysteine, PDAC

## Abstract

Resistance to immune checkpoint inhibition remains a major barrier in pancreatic cancer treatment. Here, we show that concurrent administration of probiotics restores sensitivity to anti-PD-1 therapy in pancreatic cancer mouse models. Mice treated with the combination of anti-PD-1 and probiotics demonstrate robust tumor control, accompanied by enrichment of microbial pathways governing cysteine biosynthesis, elevated serum cysteine levels, and increased T cell function. Serum cysteine levels, rather than intratumoral cysteine concentrations, inversely correlate with tumor burden. Functionally, cysteine directly promotes T cell survival, activation, and cytotoxicity while its restriction induces uncoupled transcriptional-translational stress and impairs T cell function. Oral cysteine supplementation synergizes with anti-PD-1 therapy in pancreatic cancer mice, reducing tumor burden and enhancing intratumoral T cell activation, phenocopying probiotics-mediated immune restoration. These findings suggest systemic cysteine availability as a tractable metabolic target to enhance cancer immunotherapy.

## INTRODUCTION

Immune checkpoint blockade has transformed cancer therapy, yet its efficacy remains limited in immunologically cold tumors such as pancreatic ductal adenocarcinoma (PDAC)^1,2^. Despite the presence of tumor-reactive T cells, PDAC exhibits profound resistance to immune checkpoint inhibitors (ICIs), underscoring the presence of dominant suppressive mechanisms that constrain antitumor immunity^3^. While tumor⍰intrinsic oncogenic pathways and dense stroma have long been implicated in immunotherapy resistance, multiple lines of evidence now suggest a regulatory role of systemic metabolic and microbial cues. Indeed, multiple gut microbiome⍰derived metabolites are found to modulate T cell function and immune checkpoint responsiveness beyond the local tumor microenvironment^4–6^.

The gut microbiome has emerged as a key modulator of antitumor immunity and therapeutic response to immune checkpoint blockade across multiple cancer types. Specific gut bacterial taxa and microbial functional pathways have been correlated with improved clinical outcomes in patients receiving ICIs. Gut bacteria genera such as *Eubacterium*, *Lactobacillus*, and *Streptococcus* were positively associated with anti-PD-1/PD-L1 response across different gastrointestinal (GI) cancer types^7^. For example, *Lactobacillus reuteri* protects against GI cancers by enhancing innate and adaptive immunity^8–10^. In non-small cell lung cancer patients, responders to ICI have a distinct microbial community structure, with significantly enriched taxa in the genera *Ruminococcus*, *Akkermansia*, and *Faecalibacterium* compared with non-responders^11^. Alongside this, enriched metabolic pathways related to nucleotide biosynthesis and short⍰chain fatty acid production are observed^7,11^. However, the causal mechanisms linking gut microbial composition to ICI efficacy remain incompletely defined, particularly in PDAC. Elucidating how probiotic interventions modulate cancer-associated dysbiosis and generate beneficial microbial metabolites that directly or indirectly overcome the profound immunosuppression and therapeutic resistance characteristic of PDAC represents an important and timely area of investigation.

T cell activation, survival, and effector differentiation are tightly coupled to metabolic fitness and nutrient availability. Within the tumor microenvironment, nutrient competition, oxidative stress, and limited biosynthetic capacity impose metabolic constraints that impair T cell function. Among amino acids, Cysteine (Cys) or Cystine (CySS) serves as a critical substrate for protein synthesis, redox homeostasis through glutathione production and signaling. Cys is a conditionally non-essential amino acid that can be synthesized from methionine and serine under normal physiological conditions^12,13^. In circulation, it predominantly presents as CySS, the oxidized dimer form of Cys, and is the primary form that can be imported into cells by its transporter, system xc^-14,15^. T cells lack cystathionase for *de*□*novo* Cys synthesis and are largely dependent on extracellular Cys or CySS uptake to meet their metabolic demands during activation and proliferation^16–18^. Despite the fundamental metabolic constraint, our understanding of how cysteine availability qualitatively and quantitatively shapes T cell biology is limited.

Existing evidence for cysteine-T cell biology is largely derived from *in vitro* studies using supraphysiological concentrations (10-50 mM), far exceeding the physiological and most pathological concentrations^19,20^. These studies do not define how physiologically relevant fluctuations in Cys/CySS regulate T cell activation, survival, and effector differentiation, nor do they clarify the molecular mechanism involved. A recent work demonstrates that dietary cysteine can enhance T cell-mediated functions, including CD8⁺ T cell-derived IL-22 production, which in turn regulates tissue homeostasis and immune support of intestinal stemness^21^. However, whether Cys/CySS availability serves as a limiting metabolic checkpoint that constrains the antitumor immunity or modulates the ICI efficacy *in vivo* has not been systematically investigated. Thus, significant gaps remain in our understanding of cysteine biology in T cells, particularly in tumor immunity and therapeutic contexts.

Here, we hypothesized that tumor-associated gut dysbiosis limits systemic cysteine availability, which can be restored by probiotics to preserve T cell metabolic fitness and reverse the resistance to PD-1 blockade in PDAC. Using orthotopic and autochthonous PDAC mouse models, we show that anti-PD-1 monotherapy provides minimal benefit, whereas probiotic-mediated microbiome remodeling restores therapeutic sensitivity. We identify Cys biosynthesis as a key microbial functional pathway enriched by probiotics when combined with anti-PD-1 treatment and demonstrate that systemic, rather than intratumoral, Cys concentration correlates with tumor control. Through integrated metagenomic, metabolic, transcriptomic, and functional analyses, we define Cys as a limiting systemic nutrient that governs T cell survival, transcription-translational coupling, and effector function. Finally, we show that oral supplementation of N-acetyl cysteine (NAC), a cell-permeable Cys pro-drug and an FDA-approved medication for acetaminophen toxicity^22^, phenocopies probiotic-mediated immune restoration and synergizes with anti-PD-1 therapy *in vivo*. Together, our findings uncover a microbiome-Cys-T cell axis that regulates ICI efficacy and suggests Cys as a tractable target to overcome immunotherapy resistance.

## RESULTS

### Probiotics Restore Anti-PD-1 Efficacy by Remodeling Gut Microbiota

We have found that both the pancreatic and gut microbiomes are distinct in hosts with PDAC^23^. Comparing the microbiomes of tumor-free and PDAC mice, *Lactobacillus*, *Akkermansia*, and *Ruminococcus* were associated with tumor-protection and immune activation^23^. *Akkermansia muciniphila* improves gut barrier integrity, limiting translocation^24^, and has been shown to enhance the response to anti-PD-1 therapy by promoting dendritic cell maturation and cytotoxic T cell infiltration^25^. *L. reuteri* protects against GI cancers via enhancement of innate and adaptive immunity^8–10^. Therefore, we chose these bacterial taxa as putative ‘probiotics’ to test if they can enhance the efficacy of anti-PD-1 in PDAC. We applied an orthotopic murine PDAC model by implanting the murine pancreatic cancer cell line, KPC cells, in the pancreases of 10-week-old male mice. Mice were then randomly assigned to receive isotype IgG injections (Control group), PD1-antibody injections (Anti-PD-1 group), or PD1-antibody injections combined with probiotic solution through oral gavage (Combination group) (ddH_2_O oral gavage was performed in the Control group and Anti-PD-1 group) (Figure 1A). After 6 weeks of treatment, there was no statistically significant difference in tumor size between the control and anti-PD-1 monotreatment groups (p=0.0939). In contrast, the combination group exhibited a significant reduction of tumor burden (p=0.0027 versus an anti-PD-1 group and p<0.0001 versus a Control group) (Figure 1B). Interestingly, while alpha diversity metrics remaining unchanged (Figure S1A), PCoA based on multiple beta diversity metrics including Weighted-UniFrac distances (Figure 1C), Aitchison distance (Figure S1B), and Bray-Curtis dissimilarity (Figure S1C) revealed a significant shift in microbial community structure of the fecal samples from before (baseline) and after (endpoint) treatment in the combination group. Only Aitchison distance analysis exhibited a significant difference in mice receiving anti-PD-1 monotherapy (p=0.002), whereas other beta diversity metrics did not (Figure S1B), suggesting subtle treatment-associated compositional shifts.

**Figure 1.**
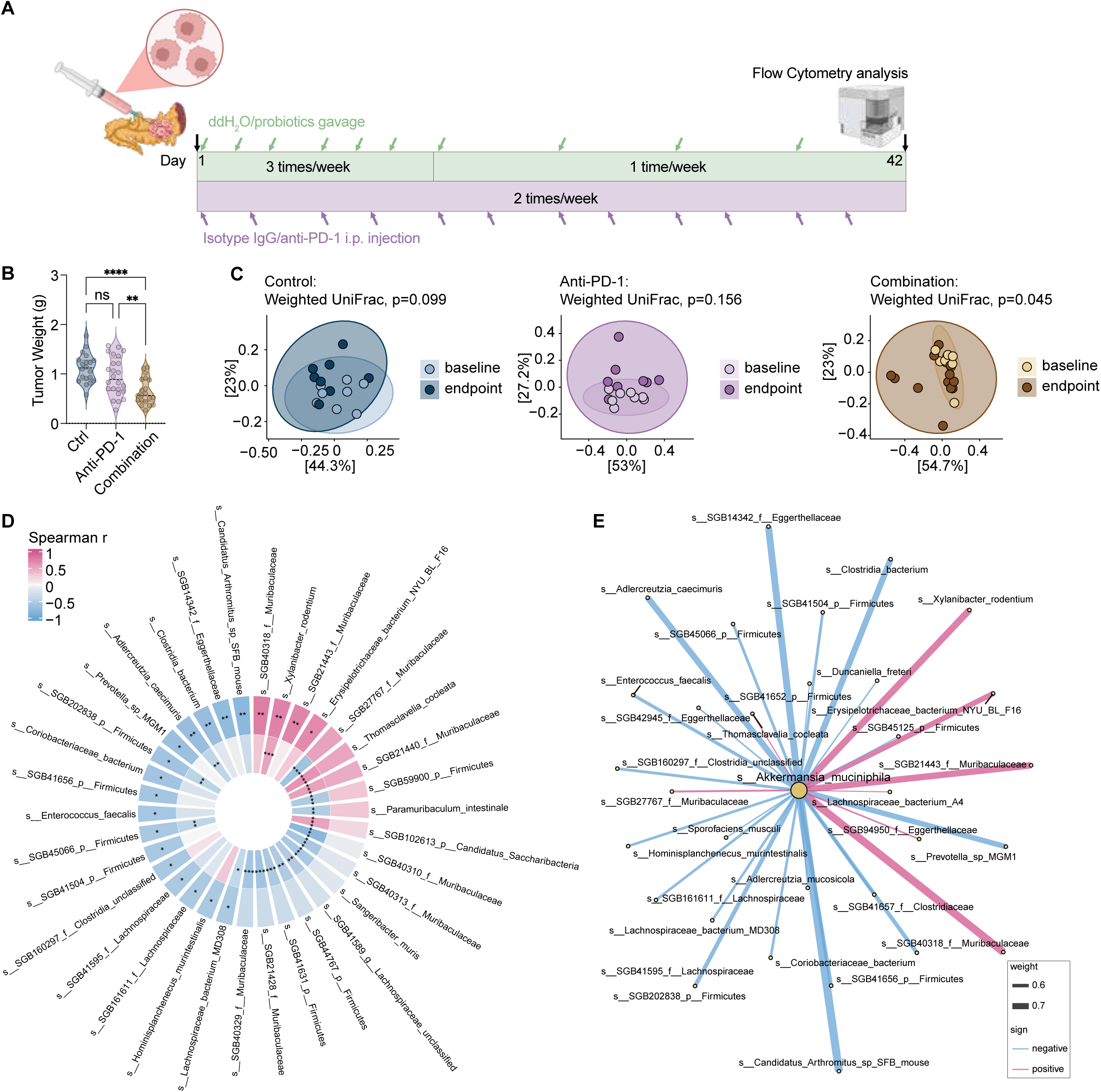
Probiotics restore sensitivity to PD-1 blockade and modulate microbiome community in PDAC mice. (A and B) Schematic (A) and tumor weight (B) of KPC tumor-bearing mice receiving isotype IgG (n=20) or anti-PD-1 injections combined with ddH_2_O (n=24) or probiotics (n=20) gavage for 6 weeks. Created with BioRender.com (A). Data are represented as mean ± SEM, whereas ns p>0.05, **p<0.01, and ****p<0.0001 by one-way ANOVA followed by Fisher’s LSD tests (B). (C) PCoA of beta diversity based on the Weighted UniFrac distance matrix, comparing baseline and endpoint sample community structure in the Ctrl, Anti-PD-1, and Combination groups (n pairs=8, 8, 13, respectively). (D) Ring heatmap of Spearman correlation coefficients between *A. muciniphila* and other species, p<0.05 and |r|>0.5 at either baseline or endpoint. The inner ring shows all baseline samples (n=37), and the outer ring shows the combination-treatment endpoint samples (n=13). (E) Spearman correlations (|r|>0.5) between *A. muciniphila* and other microbial species at the endpoint of the combination group (n=13). Edge color and thickness indicate correlation direction (red: positive; blue: negative), and strength, respectively. See also Figure S1 and Table S1.

Species-level Spearman correlations between *A. muciniphila* and other species were calculated at baseline and endpoint to evaluate shifts in microbial association patterns (Figure 1D and Table S1). At baseline, predominantly negative correlations alongside positive correlations of a few select Muribaculaceae members with *A. muciniphila* were observed (Figure 1D and Table S1). After the combination treatment, a shift in the correlation landscape is presented by strengthened correlations with *Candidatus Arthromitus* (SFB), *Lachnospiraceae bacterium* MD308, *Xylanibacter rodentium* (formerly *Prevotella rodentium*), and *Erysipelotrichaceae bacterium*. New correlations with Eggerthellaceae members, more Muribaculaceae members, Firmicutes members, and multiple Lachnospiraceae SGBs appeared at the endpoint (Figures 1D-1E and Table S1). Muribaculaceae contribute to propionate, acetate, and butyrate production in the gut^26,27^, and *X. rodentium* possesses a diversity of genes for carbohydrate-active enzymes^28^, suggesting a modulation in complex carbohydrate metabolism with cooperative cross-feeding interactions after the combination treatment.

### Probiotics Boost Intratumoral T Cell Function

We next characterized the intratumoral immune environment by flow cytometry and observed increased infiltration of CD45^+^ leukocytes, CD4^+^, and CD8^+^ T cells (Figures 2A-2C). Importantly, the activated CD4^+^ and CD8^+^ T cells increased in the PDAC tumors from the combination group (Figures 2D-2E). Enhanced CD4^+^ T cell infiltration and reduced granulocyte infiltration were further validated by immunofluorescence, with quantification performed using the HALO platform (Figures 2F-2H). We also validated the efficacy of the combination therapy in a widely used genetically engineered PDAC mouse model, KC mice, by crossing p48^Cre^ and LSL-KRAS^G12D^ mice as described previously^23^ (Figure 2I). The pancreata of KC mice express oncogenic *Kras* in acinar cells, display the highest PanINs proliferative index, and develop more than 80% neoplastic ducts by 7 to10 months^29^, mimicking the stepwise progression of human PDAC. KC mice in the combination treatment group had significantly lower tumor weight than the control group and anti-PD-1 monotherapy group (Figure 2J), similar to the observations in the KPC orthotopic PDAC model. Moreover, combination-treated KC mice exhibited increased intratumoral infiltration of CD45^+^ leukocytes, CD4^+^ T cells, CD8^+^ T cells, as well as the activated CD4^+^ and CD8^+^ T cells, as we observed in the KPC model (Figures 2K-2O).

**Figure 2.**
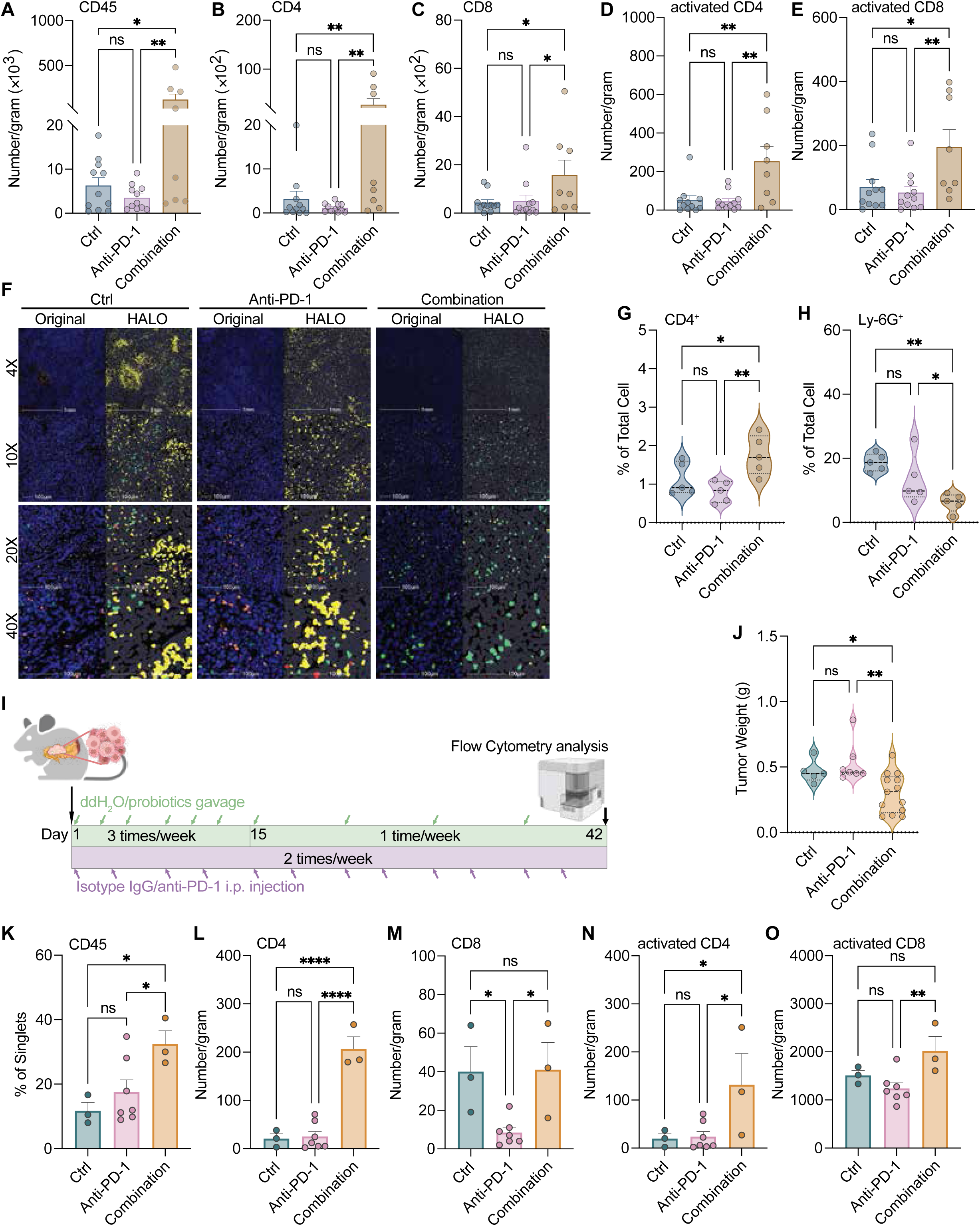
The combination of anti-PD-1 and probiotics boosts T-cell infiltration in PDAC mouse models. (A-E) Tumor-infiltrated CD45^+^ (A), CD4^+^ (B), CD8^+^ (C), activated CD4^+^ (D), and CD8^+^ (E) (define as CD25 or/and CD69 positive) T cells detected by flow cytometry from KPC tumor-bearing mice receiving isotype IgG (n=11) or anti-PD-1 injections combined with ddH_2_O (n=11) or probiotics (n=8) gavage for 6 weeks. (F)) Representative original immunofluorescence and HALO analysis images of tumor sections staining with CD4 (in green), Ly-6G (in yellow), and FOXP3 (in red). (G and H) Tumor-infiltrated CD4^+^ (G) and Ly-6G^+^ (H) cells quantified by HALO platform (n=5 per group). (I) Schematic of treatments in KC transgenic PDAC mice. Created with BioRender.com. (J) Tumor weight of KC transgenic mice receiving isotype IgG (n=5) or anti-PD-1 injections combined with ddH_2_O (n=7) or probiotics (n=13) gavage for 6 weeks. (K-O) Tumor-infiltrated CD45^+^ (K), CD4^+^ (L), CD8^+^ (M), activated CD4^+^ (N), and CD8^+^ (O) (defined as CD25 or/and CD69 positive) T cells detected by flow cytometry from KC mice receiving isotype IgG (n=3) or anti-PD-1 injections combined with ddH_2_O (n=7) or probiotics (n=3) gavage for 6 weeks. Data are represented as mean ± SEM, whereas ns p>0.05, *p<0.05, **p<0.01, and ****p<0.0001 by one-way ANOVA followed by Fisher’s LSD tests. See also Figure S2.

Accumulating evidence indicates that the gut microbiome contributes to immune modulation induced by ICIs^30,31^. Nevertheless, it remains unclear whether antibiotic-mediated microbial depletion interferes with or enhances the probiotic-induced reprogramming of host immunity that promotes sensitivity to anti-PD-1 therapy. To directly address this question, we employed a well-characterized bacterial flora removal protocol, i.e., oral antibiotic treatment^32,33^. Antibiotics were administered two weeks prior to KPC tumor implantation, followed by anti-PD-1 injections with ddH_2_O or probiotics gavage for 6 weeks (Figure S2A). Removal of native gut microbiota abolished the efficacy of combination treatment observed in mice with intact gut flora; instead, a similar tumor burden to the anti-PD-1 group or control group was observed (Figure S2B). The intratumoral lymphocytes infiltration, including HALO-quantified CD4^+^ T cells, Ly-6G^+^ granulocytes, and CD8^+^ T cells, were similar among groups (Figures S2C-S2E). These findings highlight an essential role of the microbial community in establishing probiotic-enhanced immune responses.

### Enriched Cysteine Biosynthesis Pathways in Remodeled Microbiome

The requirements of an intact microbial community for probiotic-enhanced anti-PD-1 efficacy strongly suggest the importance of integrated microbial function networks. Thus, we performed functional profiling at the pathway-level (MetaCyc) from shotgun metagenomic data to quantify the metabolic potential of the microbial community. We identified 18 MetaCyc pathways that were significantly altered during treatment between anti-PD-1 monotherapy and combination groups (Table S2). Two direct Cys biosynthesis pathways, PWY-6293 and PWY-801, as well as two pathways related to Cys metabolism, were significantly altered in the combination group (Figure 3A).

**Figure 3.**
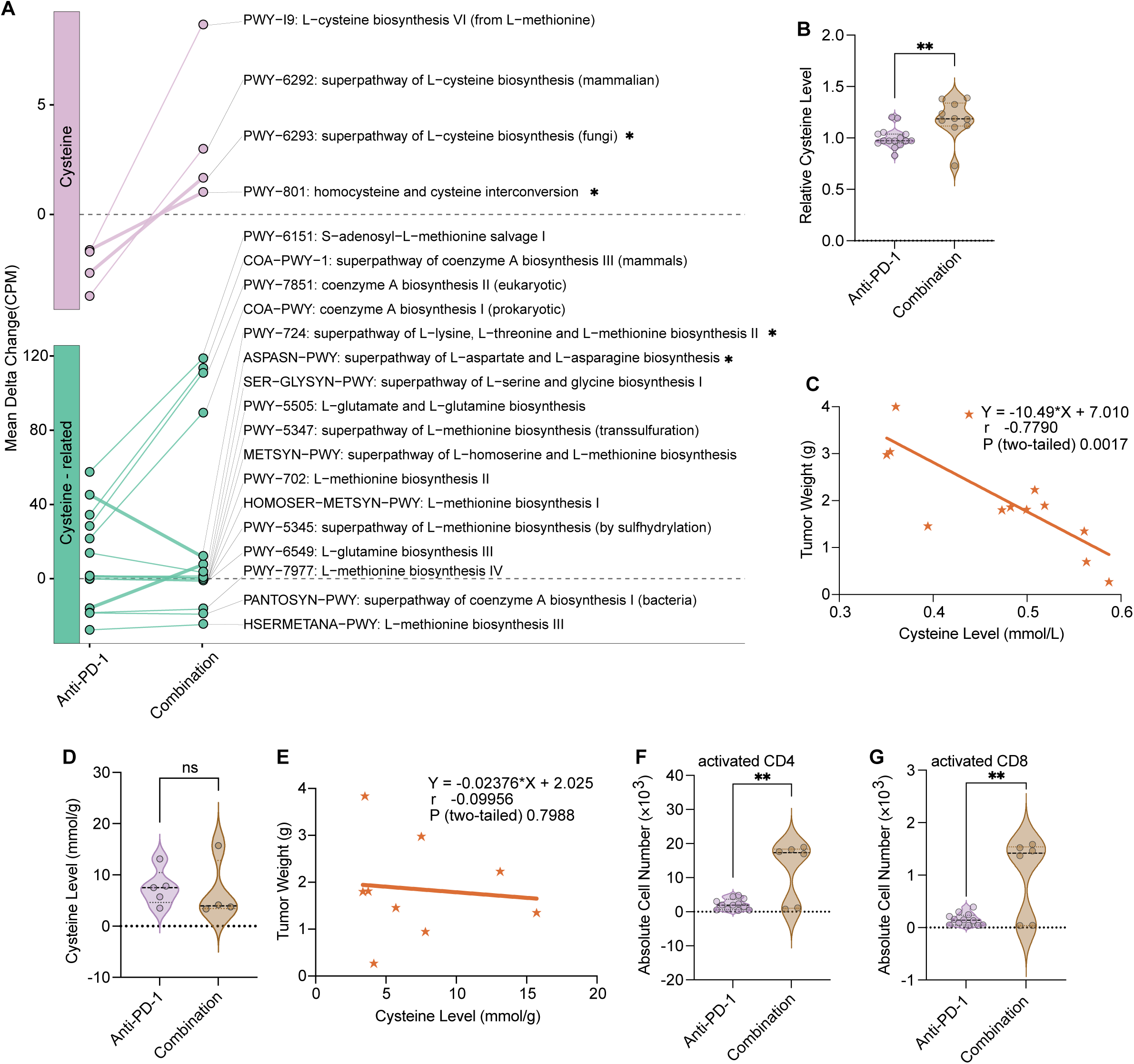
Probiotic-driven microbiome remodeling enriches cysteine biosynthetic pathways. **(A)** Line plot connecting group-mean delta copies per million (CPM) for cysteine-related pathways between anti-PD-1(n=8) and combination(n=13) groups. Statistical significance is indicated by thicker line weights and asterisks, whereas *p<0.05. (B) Relative serum cysteine levels of the KPC tumor-bearing mice receiving anti-PD-1 (n=15) or combination (n=10) treatment for 6 weeks. (C) Correlation between serum cysteine levels and tumor weights (n=13). The Pearson correlation coefficient was used to assess the strength and direction of the relationship. (D) Tumor tissue cysteine levels of the KPC tumor-bearing mice receiving anti-PD-1 (n=5) or combination (n=4) treatment for 6 weeks. (E) Correlation between KPC tumor tissue cysteine levels and tumor weights (n=9). The Pearson correlation coefficient was used to assess the strength and direction of the relationship. (F and G) The number of activated CD4^+^ (F) and CD8^+^ (G) (defined as CD25 or/and CD69 positive) T cells in mesenteric lymph nodes were detected by flow cytometry from KPC tumor-bearing mice receiving anti-PD-1 (n=12) or combination (n=6) treatment for 6 weeks. Data are represented as mean ± SEM, whereas ns p>0.05 and **p<0.01 by unpaired two-tailed Student’s t-tests (B, D, F, and G). See also Table S2.

We next measured the serum Cys levels of the KPC tumor-bearing mice and found significantly higher Cys concentrations in the combination group than the anti-PD-1 group (Figure 3B). Importantly, serum Cys levels significantly and inversely correlated with tumor weight (r=-0.7790, p=0.0017; Figure 3C). In contrast, intratumoral Cys levels did not differ between groups (Figure 3D) and showed no significant association with tumor burden (r=-0.09956, p=0.7988; Figure 3E). It is plausible that systemic Cys availability, rather than intratumoral Cys concentrations, influence the tumor burden, and this possibility led us to examine whether T cell activation occurs in secondary lymphoid organs rather than at the tumor site. Indeed, the number of activated CD4^+^ and CD8^+^ T cells in the mesenteric lymph nodes was significantly increased in the combination-treated group (Figures 3F-3G), which indicates that circulating Cys directly primes T cells prior to tumor infiltration, thereby promoting antitumor immunity.

### Cyst(e)ine Promotes T Cell Survival, Activation, and Cytotoxic Function

Cys is present in the extracellular space primarily in its oxidized form, CySS, which is the primary form taken up by cells^34^. This uptake occurs mainly through the system xc^-^ transporter, composed of SLC7A11 and SLC3A2^14,15^. Following cellular uptake, CySS is rapidly reduced to Cys in the cytosol, contributing to intracellular redox balance and glutathione synthesis^13,35^. The presence or absence of the main transporter and other known transporters capable of transporting Cys/CySS were inferred in KPC and T cells by semi-quantitative PCR (Table S3). We then compared their mRNA expression levels in KPC cells, unactivated splenic T cells (unactivated T cells), and anti-CD3 plus anti-CD28 activated T cells (activated T cells). The mRNA levels of genes encoding for system xc^-^ (Slc7a11 and Slc3a2; Figures S3A) and two genes of Cys transporters Slc1a4 and Slc1a5 (Figures S3B) were expressed at significantly higher levels in the KPC cells than in unactivated T cells. The markedly lower expression of Slc7a11 and Slc3a2 in unactivated T cells strongly suggested that they are unable to effectively compete with KPC tumor cells for CySS. Thus, unactivated T cells are metabolically disadvantaged in accessing extracellular CySS within the tumor microenvironment.

Notably, the mRNA levels of all these transporters were significantly upregulated in activated T cells (Figures S3A-S3B). However, the mRNA level of Slc7a11, which encodes the catalytic transporter subunit of system xc^-^, remained lower in activated T cells than KPC cells, despite the similar expression levels of Slc3a2, which encodes a chaperone that stabilizes the transporter complex at the plasma membrane (Figure S3A). We conducted dose-responsive CCK-8 proliferation assays to determine the half-maximal effective concentration (EC_50_) of Cys and CySS in all three cell types. The cell-permeable Cys precursor, N-acetylcysteine (NAC), was added at the indicated concentrations to the culture medium containing a physiological concentration of CySS (20 µM, comparable to levels in mouse serum) (Figure S3C). Activated T cells exhibited markedly enhanced responsiveness to Cys, as indicated by a substantial decrease in EC_50_ values (from 1276 µM in unactivated cells to 234.1 after activation) (Figure S3C and Table S4). Together with the up-regulated expression of transporters, these results indicated that activated T cells are more sensitive to CySS and Cys and exhibit increased dependency on this amino acid upon activation. Importantly, in the presence of physiological concentration of CySS, the EC_50_ value for additional Cys in activated T cells was approximately two-fold lower than that observed in KPC tumor cells (489.0 µM) (Table S4 and Figure S3C). Thus, augmenting Cys availability beyond physiological CySS levels may confer a competitive advantage to activated T cells over KPC cells, supporting the use of Cys supplement to elevate systemic Cys levels and enhance T cell function.

To reveal the dependency of Cys/CySS on the growth of unactivated T, activated T, and KPC cells, graded concentrations of Cys (Figure S3D) or CySS (Figure S3E) were added to the culture medium free of Cys/CySS. The EC_50_ values of Cys and CySS in activated T cells were significantly lower than those in unactivated T cells, representing a reduction of 76% and 62%, respectively (Table S4). When transporter-independent Cys was provided in the form of NAC, similar EC_50_ values were noted in the activated T cells (127.1 µM) and KPC cells (124.0 µM) (Figure S3D). In contrast, KPC cells showed a marked advantage over T cells when transporter-dependent CySS uptake was the sole available source (Figure S3E), consistent with their higher expression of transporters. These results strongly indicate that T cells are at a disadvantage when competing with KPC cells for CySS within the tumor.

To test whether Cys supplementation directly modulates T cell metabolic fitness and improves their function, murine splenic T cells were cultured in medium containing 20 µM CySS, with or without Cys supplementation. Supplemental Cys significantly enhanced T cell growth and survival, independent of anti-CD3/anti-CD28 stimulation (Figures 4A-4F). Cys supplementation also promoted T cell activation (Figures 4G-4I) and favored differentiation toward a central memory phenotype, defined by CD44⁺CD62L⁺ double-positive expression (Figures 4G and 4J–4K). In parallel, PD-1 expression increased on both CD4⁺ and CD8⁺ T cells (Figures 4G and 4L–4M), accompanied by elevated TNF-α and IFN-γ production in CD8⁺ T cells (Figures 4G and 4N–4O), supporting enhanced effector functionality rather than exhaustion.

**Figure 4.**
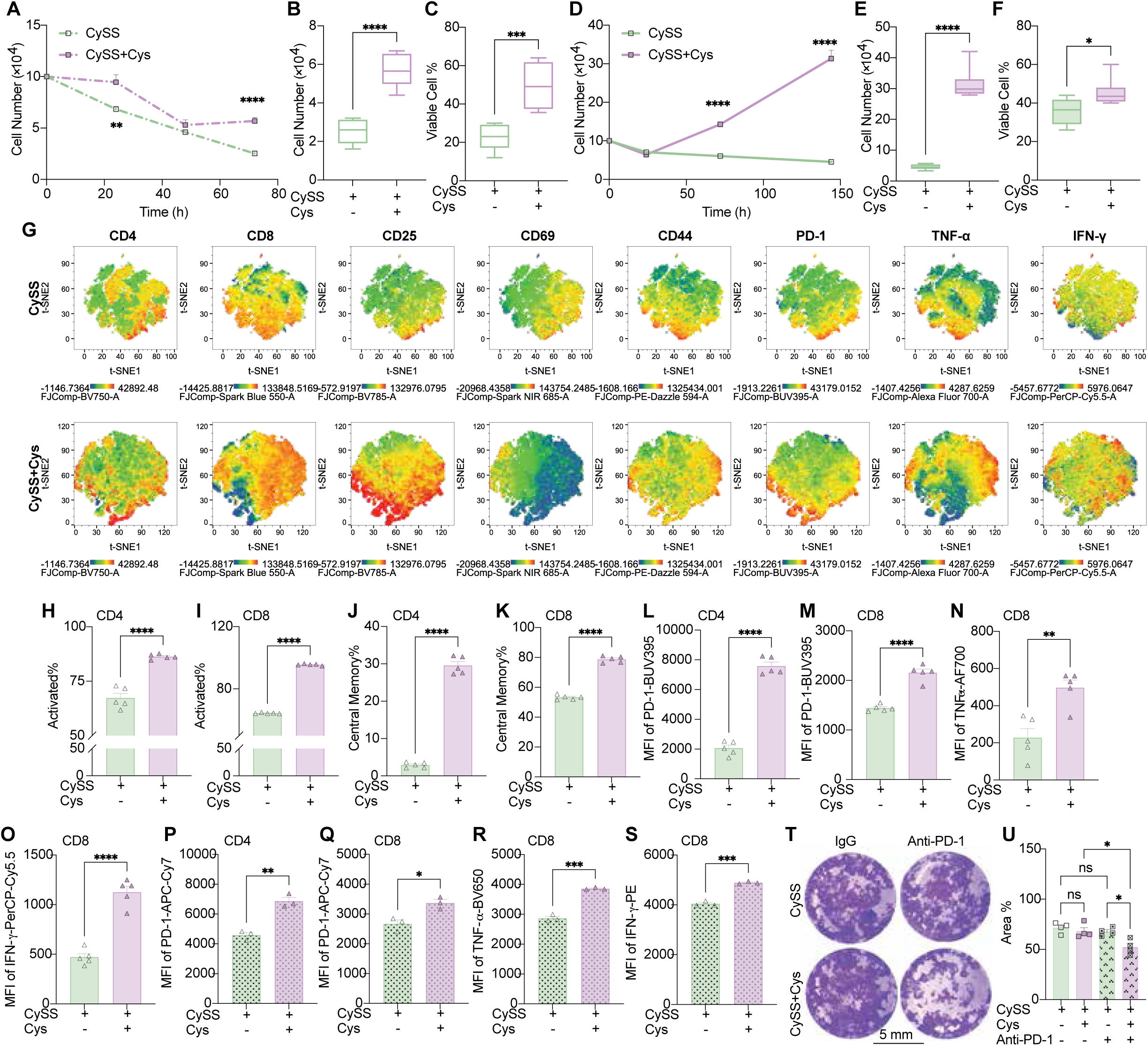
Cysteine supplementation enhances T cell survival, activation, and cytotoxicity *in vitro (A)* Murine splenic T cell growth curve. (B and C) Absolute number per well (B) and viability (C) of murine splenic unactivated T cells after 3 days of culturing. (D) Murine splenic T cell growth curve with anti-CD3 and anti-CD28 activation. (E and F) Absolute number per well (E) and viability (F) of murine splenic T cells following 6 days of anti-CD3 and anti-CD28 activation. (G) Representative T-distributed Stochastic Neighbor Embedding (t-SNE) plots of 6 days-activated murine splenic T cells. (H-K) The proportions of activated subsets (defined as CD25 or/and CD69 positive) (H and I) and central memory subsets (defined as CD44 and CD69 double-positive) (J and K) of murine splenic activated T cells. (L and M) Mean fluorescence intensity (MFI) of Brilliant Ultraviolet 395-labelled PD-1 on murine splenic activated CD4^+^ (L) and CD8^+^ (M) T cells. (N and O) MFI of Alexa Fluor 700-labelled TNF-□ (N) and PerCP-Cy5.5-labelled IFN-γ (O) on murine splenic activated CD8+ T cells. (P and Q) MFI of APC-Cyanine 7-labelled PD-1 on human CD4^+^ (P) and CD8^+^ (Q) T cells activated by human T-activator Dynabeads™ for 2 days. (R and S) MFI of Brilliant Violet 650-labelled TNF-□ (R) and PE-labelled IFN-γ (S) expression on human CD8^+^ T cells activated by human T-activator Dynabeads™ for 2 days. (T) Representative photos of KPC cells stained with crystal violet after co-culturing with activated T cells for 48 hours. (U) Quantification of crystal violet-stained area from panel T using QuPath v.0.6.0. n=6 per group for panels A-F. n=5 and 4 per group for panels H-O and U, respectively. N=3 per group for panels P-S. Data are represented as mean ± SEM except for panels G and T. Unpaired two-tailed Student’s t-tests were used except for panels A, D, G, T, and U. Multiple unpaired t-test corrected by the Holm-Šidák method was used for panels A and D, while One-way ANOVA followed by Tukey’s post hoc tests for panel U. Statistical significance was defined as follows: ns p>0.05, *p<0.05, **p<0.01, ***p<0.001, and ****p<0.0001. See also Figures S3-S7 and Table S3-S4.

To assess whether these findings are broadly applicable, we extended our analyses to multiple human T cell systems. Similar to murine T cells, human peripheral blood T cells cultured with supplemental Cys exhibited increased PD-1 expression in both CD4⁺ (Figure 4P) and CD8⁺ T cells (Figure 4Q), along with enhanced production of TNF-α (Figure 4R) and IFN-γ (Figure 4S). Notably, genetically engineered chimeric antigen receptor (CAR) T cells also displayed augmented TNF-α and IFN-γ production in response to Cys supplementation (Figures S4A-S4I). Collectively, these results demonstrate that Cys supplementation robustly enhances the survival, activation, and effector function in murine and human T cells *in vitro*.

We next evaluated whether Cys-enhanced T cell viability and activation lead to improved cytotoxic function. Activated T cells preconditioned with or without supplemental Cys were co-cultured with KPC cells at a 2:1 effector to target ratio for 48 hours. While Cys supplementation alone did not significantly augment T cell killing of KPC cells, Cys-preconditioned T cells exhibited markedly enhanced cytotoxicity when combined with PD-1 antibody treatment (Figures 4T-4U).

### Cystine Deprivation Impairs T Cell Viability and Activation but Is Reversed by Cell-Permeable Cysteine

We further examined the requirement of extracellular CySS to support T cell survival and activation by completely eliminating CySS from the culture medium. As anticipated, CySS deprivation markedly impaired the growth of both unactivated and activated T cells and significantly reduced their survival (Figures S5A-S5F). Consistent with these effects, activated T cell frequencies and central memory T cell populations decreased under CySS-deficient conditions, which was accompanied by reduced PD-1 expression and decreased TNF-α production (Figures S5G-S5N).

We then evaluated whether the functional impairments induced by CySS deprivation could be rescued by supplementation of Cys. Notably, adding Cys effectively restored T cell growth and survival (Figures S6A-S6F), increased the proportions of activated and central memory T cell subsets (Figures S6G-S6K), and reinstated PD-1 expression on CD4^+^ and CD8^+^ cells (Figures S6G and S6L-6M), accompanied by increased TNF-α production in CD8^+^ cells (Figures S6G and S6N). These findings further verified the vital function of Cys/CySS during T cell activation.

As a key precursor of glutathione synthesis^36^, cysteine is central to cellular redox homeostasis and protection against oxidative stress and ferroptosis. We therefore assessed lipid peroxidation and intracellular reactive oxygen species (ROS) levels using Liperfluo and CellROX™ Deep Red staining, respectively. Under conditions of adequate Cys/CySS availability, either CySS alone, Cys alone, or supplemental Cys in the presence of physiological CySS, T cells consistently exhibited increased levels of lipid peroxidation (Figures S7A-S7G) and ROS (Figures S7H-S7N). Notably, these elevations coincided with enhanced T cell survival and activation, suggesting that metabolically fit T cells tolerate or actively engage heightened redox activity, rather than experiencing oxidative damage.

### Cysteine Supplementation Drives Proliferative and Translational Programs in T cells

To unbiasedly characterize the transcriptional programs associated with Cys-mediated enhancement of T cell survival and function, we performed bulk RNA Sequencing (RNA-seq) on antibody-activated T cells cultured at physiological CySS level, with PBS or Cys supplementation. Gene set enrichment analysis (GSEA) identified 184 positively enriched pathways and 249 negatively enriched pathways, belonging to 19 and 18 event hierarchies, respectively, following Cys supplementation (FDR-q<0.3) (Figure 5A and Table S5). Among the positively enriched pathways, 53 cell cycle-related pathways (28.8% of the total) and 41 DNA replication and repair-related pathways (22.3%) were significantly enriched in Cys supplement group. In contrast, only 4 cell cycle-related pathways (1.8%) and no DNA replication and repair-related pathways (0%) were identified among the significantly negatively enriched pathways (Figures 5A-5B and Table S5). Additionally, pathways associated with necrosis, apoptosis, and programmed cell death were strongly negatively enriched under the Cys-supplemented conditions (Figure 5C). Moreover, numerous genes involved in cell cycle regulation were upregulated in the Cys-supplemented group (Figure 5D), including Ki67, Ccna2, Ccnb1, and Ccne2, which were independently validated by qPCR at multiple time points (Figures 5E-5H). These results provide a molecular basis for the observed increases in T cell proliferation and survival *in vitro*. Furthermore, active DNA replication was confirmed using an EdU Click-iT assay, which revealed a significant increase in EdU incorporation in Cys-supplemented T cells as early as one hour after stimulation (Figure 5I).

**Figure 5.**
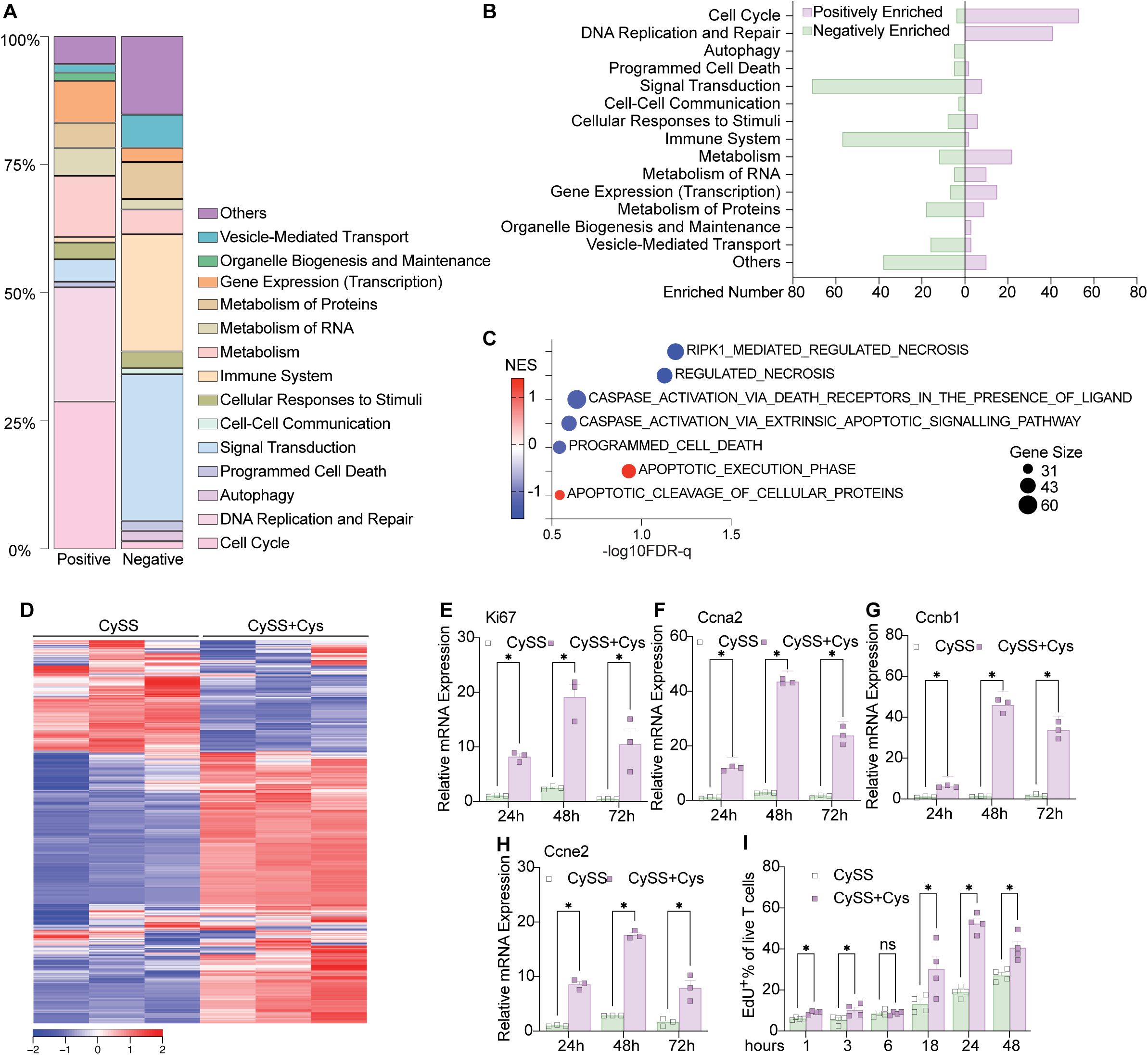
Cysteine upregulates cell cycle-related pathways and genes. (A and B) Percentages (A) and numbers (B) of positively and negatively enriched pathways in each hierarchy following Cys-supplementation analyzed by Gene Set Enrichment Analysis (GSEA) based on the M2:CP:REACTOME gene set. (C) Positively and negatively enriched pathways in the apoptosis category following Cys-supplementation. Normalized Enrichment Score (NES), significance, number of hit genes were shown by color, - log10(FDR-q) value, and bobble size, respectively. (D) Heatmap of the cell cycle-related gene expressions quantified by RNA-seq (Z-score). (E-H) Relative mRNA expression levels of Ki67 (E), Ccna2 (F), Ccnb1 (G), and Ccne2 (H) quantified by qPCR (n=3 per group). (I) DNA replication detected by the Click-iT™ EdU Cell Proliferation Kit (n=4 per group). Data are represented as mean ± SEM, whereas ns p>0.05 and *p<0.05 by multiple unpaired t-tests corrected by the Holm-Šidák method (E-I). See also Table S5.

Unexpectedly, despite their enhanced cytotoxic function, Cys-supplemented T cells exhibited significant negative enrichment across 71 signal transduction-related pathways (28.5%) and 57 immune system-related pathways (22.9%) (Figure 5A and Table S5). Classical T cell activation pathways, including 21 receptor tyrosine kinase signaling pathways and 10 Toll-like receptor cascades, were negatively enriched (Figures 6A-6B). The observed transcriptional repression contrasted with the increased T cell activation-associated proteins, including PD-1, TNF-α, and IFN-γ as demonstrated by flow cytometric analysis (Figures 4L-4O). We therefore evaluated the protein expression levels of several representative targets related to T cell function by flow cytometry and found a consistent dissociation between mRNA abundance and protein output (Figure 6C). We then directly examined whether Cys availability regulates transcriptional and translational capacity by quantifying nascent mRNA synthesis using EU incorporation and protein synthesis using puromycin incorporation in activated T cells cultured with or without supplemental Cys. Interestingly, Cys supplementation significantly enhanced both mRNA and protein synthesis (Figures 6D-6E). Given that the IFN and TNF signaling pathways were significantly negatively enriched in the Cys-supplemented group based on the RNA-seq analysis (Figure 6B), while IFN-γ and TNF-α production were markedly increased *in vitro* (Figures 4N-4O), we selected Ifng and Tnfa to further assess mRNA accumulation in stimulated T cells at different time-points. Ifng expression was significantly elevated in Cys-supplemented T cells during the early phase of stimulation (10 hours) but declined to levels below those observed in control T cells upon prolonged stimulation (24 hours) (Figure 6F). Tnfa expressions displayed a similar pattern, albeit with differences in kinetics and magnitude (Figure 6G). Notably, despite the late transcriptional decline, IFN-γ (Figure 6H) and TNF-α (Figure 6I) protein levels significantly increased in Cys-supplemented T cells at later time points.

**Figure 6.**
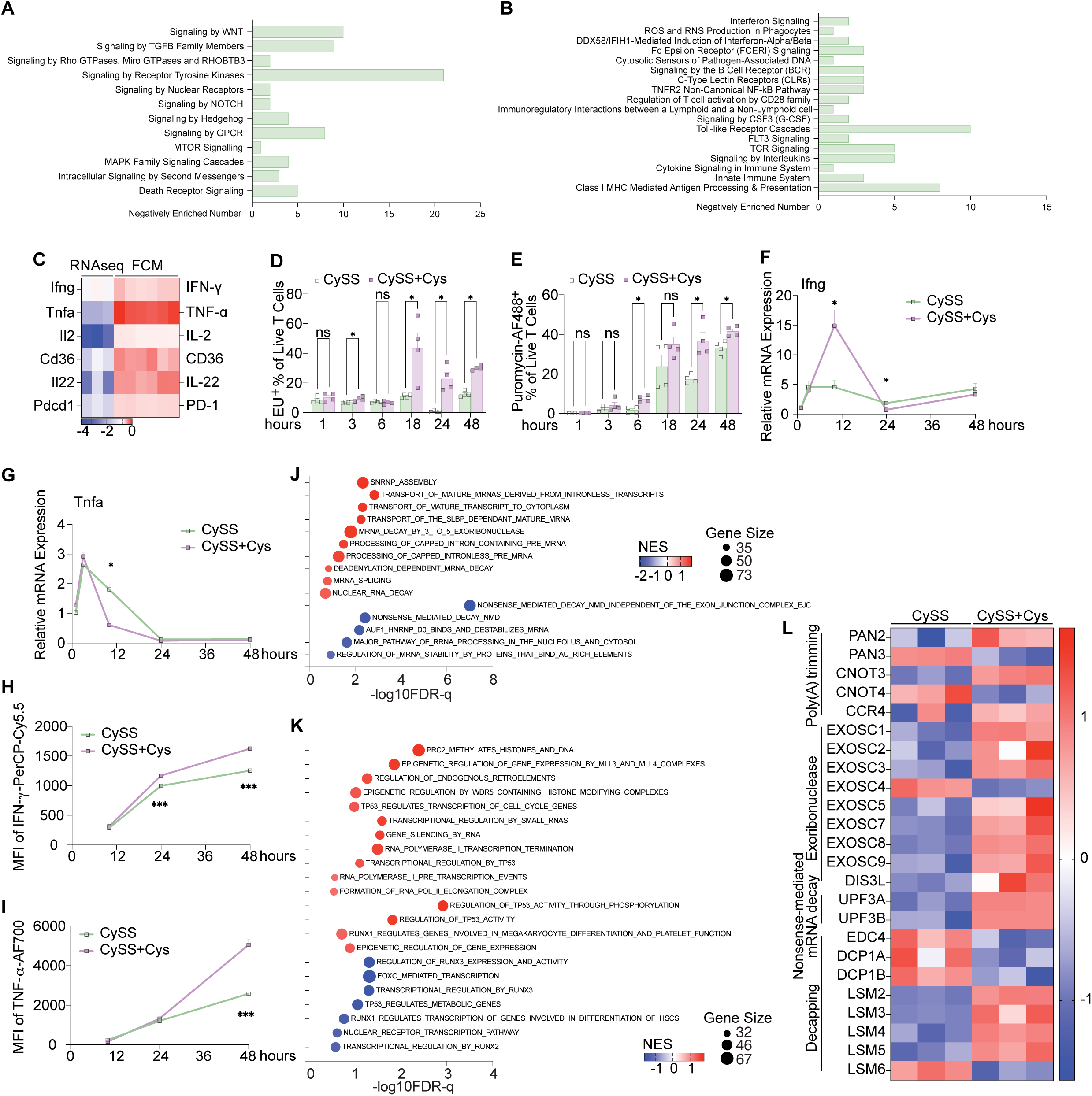
Cysteine facilitates T-cell transcription and RNA-metabolic programs. (A and B) Negatively enriched pathways in the hierarchies of signal transduction (A) and immune system (B) by additional Cys treatment. (C) Heatmap of Log2(fold change) of representative targets in mRNA levels and protein levels detected by RNA-seq (n=3) and flow cytometry (n=6), respectively. (D and E) RNA transcription (D) and protein translation (E) efficiency analyzed by EU labelling and anti-Puromycin detection (n=4 per group). (F-I) IFN-γ and TNF-□ expressions in mRNA level (F and G) and protein level (H and I) (n=3 per group). (J and K) Positively and negatively enriched pathways in the gene expression (transcription) (J) and metabolism of RNA categories (K) by additional Cys treatment. Normalized Enrichment Score (NES), significance, number of hit genes were shown by color, -log10(FDR-q) value, and bobble size, respectively. (L) Heatmap of RNA degradation-related genes quantified by RNA-seq (Z-score). Data are represented as mean ± SEM, whereas ns p>0.05, *p<0.05, and ***p<0.001 by multiple unpaired t-tests corrected by the Holm-Šidák method (D-I).

We therefore hypothesized that Cys supplementation enhances translational efficiency, thereby promoting more effective mRNA processing and efficient protein synthesis in T cells. In support of this model, pathway enrichment analysis revealed that pathways related to RNA metabolism, gene expression, and transcription, including mRNA transport and decay, were positively enriched following Cys supplementation (Figures 6J-6K). Moreover, genes involved in mRNA decay showed minimal expression in T cells cultured without Cys supplementation (Figure 6L). These results suggested that, in the absence of supplemental Cys, mRNA transcripts accumulated but were not efficiently translated, consistent with the RNA-seq profiles. Together, these data demonstrate that sufficient Cys/CySS availability is required to efficiently couple transcription with protein synthesis, thereby sustaining T cell activation and effector function.

### Cysteine Supplementation Synergizes with Anti-PD-1 Therapy to Suppress Tumor Progression In Vivo

Given the robust *in vitro* evidence that T cells benefit from Cys supplementation and its pronounced effect on PD-1 expression (Figures 4L-4M), we next evaluated whether oral Cys administration could enhance anti-PD-1 efficacy in KPC mice (Figure 7A). Cys supplementation was provided by oral gavage of NAC, which systemically increased the serum Cys levels (Figure 7B) without changing the intratumoral Cys levels (Figure 7C). Remarkably, Cys supplementation combined with anti-PD-1 substantially suppressed tumor growth (Figure 7D). Consistent with our findings in probiotic-treated mice, the combination of Cys and anti-PD-1 not only enhanced intratumoral immunity in KPC tumor tissue, as evidenced by increased infiltration of CD45^+^ leukocytes (Figure 7E), total and activated CD4^+^ and CD8^+^ T cells (Figures 7F-7J), and elevated polyfunctional IFN-γ and TNF-α double positive CD8⁺ T cells (Figures 7K), but also increased activated CD4^+^ and CD8^+^ T cells in the mesenteric lymph nodes (Figures 7L-7M).

**Figure 7.**
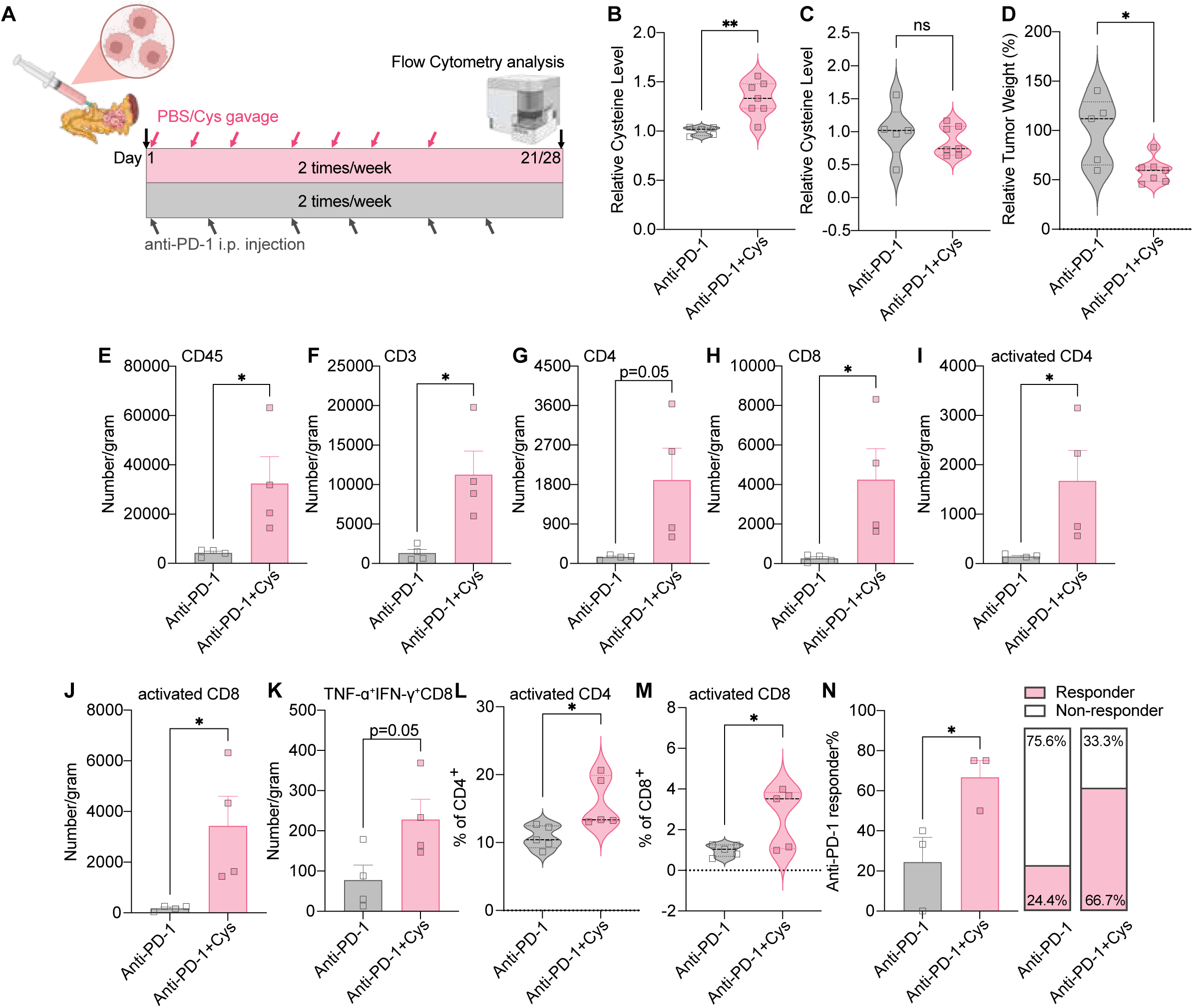
Cysteine supplementation restores anti-PD-1 efficacy in PDAC mice *in vivo*. (A) Schematic of KPC tumor-bearing mice. Created with BioRender.com. (B and C) Relative cysteine levels in serum (B) and KPC tumor tissue (C) of the KPC tumor-bearing mice receiving anti-PD-1 injections combined with PBS (n=5) or Cys (n=7) gavage for 3 or 4 weeks. (D) Relative tumor weight of the KPC tumor-bearing mice receiving anti-PD-1 injections combined with PBS (n=5) or Cys (n=7) gavage for 3 or 4 weeks. (E-J) Absolute number of tumor-infiltrated CD45^+^ (E), CD3^+^ (F), CD4^+^ (G), CD8^+^ (H), and activated CD4^+^ (I) and CD8^+^ (J) (define as CD25 or/and CD69 positive) T cells detected by flow cytometry from mice from panel A (n=4 per group). (K) Absolute number of tumor-infiltrated TNF-□ and IFN-γ double-positive CD8^+^ T cells analyzed by flow cytometry from mice from panel A (n=4 per group). (L and M) The proportions of activated CD4^+^ (L) and activated CD8^+^ (M) (defined as CD25 or/and CD69 positive) T cells in mesenteric lymph nodes detected by flow cytometry from mice from panel A (n=5 per group). (N) Anti-PD-1 responding ratio of KPC-tumor bearing mice under anti-PD-1 treatment with or without Cys gavage (each dot represents the responding ratio calculated from a single independent animal experiment). Data are represented as mean ± SEM, whereas *p<0.05 and **p<0.01 by unpaired two-tailed Student’s t-tests.

Low responsiveness to anti-PD-1 therapy represents a major barrier to effective immunotherapy in PDAC. To evaluate whether Cys supplementation improves therapeutic responsiveness in the PDAC model, anti-PD-1 responders were defined as those exhibiting at least 25% reduction in tumor weight compared with the average tumor-weight of the control group. Notably, the combination of oral Cys supplementation with anti-PD-1 treatment resulted in consistently higher responder ratios compared with anti-PD-1 monotherapy (Figure 7N). These results indicate that Cys supplementation can potentiate anti-PD-1 efficacy even in established immunotherapy-resistant tumors, highlighting a novel therapeutic strategy to overcome checkpoint blockade resistance in PDAC.

## DISCUSSION

We identified Cys availability as a key metabolic control for T cell survival, activation, and effector function. In PDAC models, facilitating T cell access to systemic Cys, either through regulation of gut microbiome composition or direct Cys supplementation orally, potentiates anti-PD-1 efficacy. Probiotic administration remodels the microbiome, increases serum Cys, and restores sensitivity to anti-PD-1 therapy in PDAC mouse models, paralleling enhanced intratumoral T cell infiltration and cytotoxicity. Community-level functional remodeling, rather than the expansion of a single species, underlies the beneficial effects of probiotics, underscoring the importance of microbiota-driven metabolic modulation for immunity.

The low expression of CySS transporters in unactivated T cells may lead to the underrated regulatory role of Cys/CySS in T cell biology. Only a few studies have explored the enhancement of T cell function using supraphysiological concentrations of Cys/CySS *in vitro*^19,20^, and the underlying mechanisms remain largely uncharacterized. Our study systematically examined the expression of a panel of CySS/Cys transporters in T cells under unactivation and activation conditions. We found that the expression of CySS transporters were markedly upregulated upon T cell activation, indicating that activated T cells exhibited substantially increased sensitivity to and dependence on Cys and CySS. Indeed, Cys directly supports T cell proliferation, survival, and differentiation into central memory phenotypes *in vitro* and promotes effector cytokine production across murine and human T cells, including engineered CAR T cells. Conversely, CySS restriction limits T cell viability, activation, and cytokine production by inducing translational stress and uncoupling mRNA transcription from protein synthesis.

Mechanistically, Cys facilitates both mRNA and protein synthesis in T cells, relieving translational arrest and enabling effective production of key activation and effector proteins, including PD-1, TNF-α, and IFN-γ. Moreover, Cys restriction reduces DNA duplication and impairs the coupling of transcription and translation. Our observation that additional Cys increases ROS levels in antibody-activated T cells is supported by previous studies^37,38^, and it was shown that antigen receptor (TcR) cross-linking rapidly induces ROS generation within minutes^36^. Increased ROS at this stage may reflect enhanced activation rather than acting as a causal determinant of T cell survival.

Prior studies have reported inconsistent and cancer-type-specific associations between serum Cys levels and cancer risk. For example, both positive^39^ and negative^40^ correlations were reported in breast cancer. CySS was listed among 4 plasma metabolites that provide a high response probability against anti-PD-1 therapy in non-small cell lung cancer^41^. These discrepancies likely reflect the context-dependent, dual role of Cys in supporting the proliferation and survival of both cancer cells and T cells. Notably, Cys has been reported to be enriched in colorectal tumors relative to matched non-tumor tissue^42^. It is plausible that microbiome-derived cysteine, by elevating systemic Cys levels, promotes T cell activation in secondary lymphoid organs, as we observed in mesenteric lymph nodes, and subsequently enhances their effector function. This effect can be further supported by the significant upregulation of PD-1 expression on T cells induced by Cys, as evidenced by the tumor-suppressive effect of the combined Cys and anti-PD-1 treatment in PDAC mice. Therefore, elevated systemic, rather than intratumoral, Cys availability may improve the efficacy of anti-PD-1 therapy in otherwise “cold” solid tumors.

Our findings highlight a Cys⍰dependent translational checkpoint that constrains T cell function under nutrient limitation. Importantly, oral Cys administration synergized with anti-PD-1 therapy to reduce KPC tumor burden and phenocopied the immunomodulatory effects of probiotics by increasing circulating Cys without elevating intratumoral Cys. These results suggest that systemic Cys availability, rather than local Cys accumulation within the tumor, can be harnessed to enhance immunotherapy in checkpoint-resistant tumors. Of note, Cys deprivation approaches, including dietary Cys restriction, inhibition of CySS transport, or systemic cysteinase administration, have suppressed tumor growth in preclinical models^43^. It is possible that a therapeutic window exists in which Cys restriction suppresses tumor growth without critically impairing T cell function. However, the double-edged nature of restricting Cys should be considered: while Cys restriction may impair tumor cell redox homeostasis and proliferation, it may simultaneously compromise anti-tumor immunity by constraining T cell survival and effector function. Ideally, Cys availability would be restricted selectively in tumor cells rather than systemically.

Our data align with growing interest in metabolic and microbial determinants of immunotherapy response. While the microbiome’s influence on checkpoint blockade with differences in taxa correlating with clinical outcomes has been described in melanoma and other cancers^44^, our findings extend this concept through systemic Cys supplementation to metabolically support T cell function and synergize with ICI in PDAC. Importantly, we also found increased PD-L1 expression in KPC cells under Cys supplement conditions (data not shown), reinforcing the rationale for combined anti-PD-1 and Cys therapy and underscoring that augmenting T cell metabolic fitness can overcome checkpoint resistance in an otherwise refractory tumor model, highlighting the potential broad applicability of Cys-assisted immune checkpoint inhibition strategies, including anti-PD-1 or anti-PD-L1 monotherapies.

In summary, this study defines a gut microbiota-systemic cystine-T cell axis that governs ICI efficacy and identifies CySS/Cys availability as a tractable metabolic target to enhance immunotherapy in immunologically resistant tumors. Additional studies are warranted to advance the translational application of these findings. The optimal strategies to modulate systemic Cys in patients, whether via diet, targeted probiotics, or supplementation, require careful clinical validation. While our findings support using NAC orally for Cys augmentation to overcome PD-1 resistance, prospective clinical studies will be necessary to confirm safety, efficacy, and biomarker correlations in human PDAC and other ICI-resistant tumors.

### Limitations of the study

The limitation of our work is the absence of a correlation analysis between systemic CySS/Cys levels in PDAC patients and the prognosis of ICI therapy. It is because CySS/Cys is not a routine parameter included in clinical blood testing. Moreover, due to its susceptibility to oxidation, the measurement requires strict antioxidant handling during sample collection and processing. Consequently, we are unable to obtain reliable CySS/Cys concentrations in cancer patients from published studies or databases.

## Supporting information

Graphical abstract

## RESOURCE AVAILABILITY

### Lead contact

Requests for further information and resources should be directed to and will be fulfilled by the lead contact, Xin Li.

### Materials availability

Materials generated in this study are available from the lead contact upon completion of a materials transfer agreement.

### Data and code availability

All data in the study are publicly available as of the date of publication.

- Shotgun metagenomic sequencing data have been deposited at NCBI Sequence Read Archive (SRA) as accession number PRJNA1432059.
- Bulk RNA Sequencing data have been deposited at NCBI SRA as accession number PRJNA1434099 and at Mendeley at DOI: 10.17632/wgzh4pzw5p.1.
- Flow cytometry raw files have been deposited at Mendeley Data at DOI: 10.17632/7bn53gnsnk.1.
- This study does not report original code.
- Any additional information required to reanalyze the data reported in this paper is available from the lead contact upon request.

## ACKNOWLEDGMENTS

We would like to thank Drs. Jie Sun, Wenbo Yan, and Jill K. Slack-Davis at UVA School of Medicine for their review of the manuscript and their helpful suggestions. The study has been partially supported by fundings DoD W81XWH-19-1-0603 to X.L., and DoD W81XWH-19-1-0605 to D.S.. We would like to acknowledge the CCSG Support by the UVA Comprehensive Cancer Center (P30 CA044579) at the Flow Cytometry Core Facility (RRID: SCR_017829) and Genome Analysis and Technology Core (RRID: SCR_018883). We thank members of the Experimental Pathology Research Laboratory (RRID: SCR_017928), which is partially supported by the New York University Langone’s Laura and Isaac Perlmutter Cancer Center Grant (P30 CA016087) and a Shared Instrumentation Grant (S10 OD021747).

## AUTHOR CONTRIBUTIONS

Conceptualization, X.W., Z.W., Y.G., and X.L.; Data Curation and Formal Analysis, X.W., Z.W., Y.G., and F.X.; Investigation and Methodology, X.W., Z.W., Y.G., F.X., J.Z., S.T., D.S., J.X., and X.L.; Project Administration, X.W., Z.W., and X.L.; Validation, X.W. and Z.W.; Visualization, X.W., Z.W., F.X. and Y.G.; Writing-original draft, X.W. and Z.W.; Writing-review & editing, X.L., X.W., Z.W., Y.G., F.X., S.T., D.S., and J.X.; Resources, X.L., D.S., and J.X.; Funding Acquisition, X.L., J.X., and D.S.; Supervision, X.L..

## DECLARATION OF INTERESTS

The authors declare no competing interests.

## SUPPLEMENTAL INFORMATION

Document S1. Tables S2–S5.

Tables S1. Spearman correlations between Akkermansia muciniphila and other species in all treatment naive baseline and combination group endpoint samples. Related to Figure 1.

## Supplemental figures

**Figure S1. Microbiome shifts in the KPC tumor-bearing mice, related to Figure 1**

(A) Alpha-diversity based on the observed SGB richness and Shannon indices between baseline and endpoint samples in the control (n=8), anti-PD-1 (n=8), and combination (n=11) groups.

(B and C) PCoA of beta-diversity based on the Aitchison distance matrix (B) and the Bray-Curtis dissimilarity matrix (C) comparing baseline and endpoint in the control (n=8), anti-PD-1 (n=8), and combination (n=13) groups.

**Figure S2. Antibiotic pretreatment negates the advantage of combination therapy, related to Figure 2**

(A) Scheme for the bacteria depletion model of KPC tumor-bearing mice. Created with BioRender.com.

(B) Tumor weight of bacteria-depleted KPC tumor-bearing mice receiving isotype IgG (n=11) or anti-PD-1 injections combined with ddH_2_O (n=12) or probiotics (n=13) gavage for 6 weeks.

(C-E) The quantification by the HALO platform of tumor-infiltrated CD4^+^ cells (C), Ly-6G^+^ cells (D), and CD8^+^ cells (E) from bacteria-depleted KPC tumors (n=6 per group, except for the probiotics group in panel E, n=5).

Data are represented as mean ± SEM, whereas ns p>0.05 by one-way ANOVA followed by Fisher’s LSD tests.

**Figure S3. Cyst(e)ine transporter expressions and the sensitivity of cyst(e)ine in KPC cells, unactivated T cells, and activated T cells, related to Figure 4**

(A and B) Relative mRNA expression of CySS transporters (A) and Cys transporters (B) in KPC cells, unactivated T cells, and activated T cells (n=3 per group). Data are represented as mean ± SEM, whereas *p<0.05, **p<0.01, and ****p<0.0001 by one-way ANOVA followed by Fisher’s LSD tests.

(C-E) CCK-8-based dose–response curves of unactivated T cells, activated T cells, and KPC cells in 20 µM CySS-based Cys treatments (C), PBS-based Cys treatments (D), and HCl-based CySS treatments (E) (n=3). Data were fitted non-linear regression analysis using a four-parameter logistic model, and the EC₅₀ values were calculated from the best-fitted curves.

See also Tables S3-S4.

**Figure S4. Cysteine supplementation enhances human GFP CD19-28ζ CAR T cells cytokine secretion, related to Figure 4**

(A) Representative flow cytometric histograms of human CD4^+^ and CD8^+^ T cells TNF-□ expression on Brilliant Violet 650 channel.

(B and C) Mean fluorescence intensity (MFI) of Brilliant Violet 650-labelled TNF- □ on human CD4^+^ (B) and CD8^+^ (C) T cells activated by human T-activator Dynabeads™ for 2 days.

(D) Representative flow cytometric histograms of human CD4^+^ and CD8^+^ T cells IFN-γ expression on PE channel.

(E and F) MFI of PE-labelled IFN-γ on human CD4^+^ (E) and CD8^+^ (F) T cells activated by human T-activator Dynabeads™ for 2 days.

(G) Representative flow cytometric contour plots of TNF- □ and IFN-γ on human CD4^+^ and CD8^+^ T cells.

(H and I) Percentage of TNF- □ and IFN-γ double-positive subset of human CD4^+^ (H) and CD8^+^ (I) T cells activated by human T-activator Dynabeads™ for 2 days.

N=3 per group. Data are represented as mean ± SEM, whereas *p<0.05, **p<0.01, ***p<0.001, and ****p<0.0001 by unpaired two-tailed Student’s t-tests except for panels A, D, and G.

**Figure S5. Deprivation of cystine dampens T-cell survival and activation *in vitro*, related to Figure 4**

A. Murine splenic T cell growth curve.

(B and C) Absolute number per well (B) and viability (C) of murine splenic unactivated T cells after 3 days of culturing.

Murine splenic T cell growth curve with anti-CD3 and anti-CD28 activation.

(E and F) Absolute number per well (E) and viability (F) of murine splenic T cells following 3 days of anti-CD3 and anti-CD28 activation.

Representative T-distributed Stochastic Neighbor Embedding (t-SNE) plots of 6-day-activated murine splenic T cells.

(H-K) The proportions of activated subsets (defined as CD25 or/and CD69 positive) (H and I) and central memory subsets (defined as CD44 and CD69 double-positive) (J and K) of murine splenic activated T cells.

(L and M) Mean fluorescence intensity (MFI) of Brilliant Ultraviolet 395-labelled PD-1 on murine splenic CD4^+^ (L) and CD8^+^ (M) T cells.

MFI of Alexa Fluor 700-labelled TNF- □ on murine splenic CD8^+^ T cells.

n=12 per group for panels A-F. n=6 per group for panels H-N. Data are represented as mean ± SEM except for panel G. Unpaired two-tailed Student’s t-tests were used except for panels A, D, and G. Multiple unpaired t-test corrected by the Holm-Šidák method was used for panels A and D. Statistical significance was defined as follows: *p<0.05, **p<0.01, ***p<0.001, and ****p<0.0001.

**Figure S6. Cysteine rescues T-cell survival and activation from cystine deprivation in vitro, related to Figure 4**

A. Murine splenic T cell growth curve.

(B and C) Absolute number per well (B) and viability (C) of murine splenic unactivated T cells after 2 days of culturing.

Murine splenic T cell growth curve with anti-CD3 and anti-CD28 activation.

(E and F) Absolute number per well (E) and viability (F) of murine splenic T cells following 3 days of anti-CD3 and anti-CD28 activation.

Representative T-distributed Stochastic Neighbor Embedding (t-SNE) plots of 6-day-activated murine splenic T cells.

(H-K) The proportions of activated subsets (defined as CD25 or/and CD69 positive) (H and I) and central memory subsets (defined as CD44 and CD69 double-positive) (J and K) of murine splenic activated T cells.

(L and M) Mean fluorescence intensity (MFI) of Brilliant Ultraviolet 395-labelled PD-1 on murine splenic activated CD4^+^ (L) and CD8^+^ (M) T cells.

MFI of Alexa Fluor 700-labelled TNF- □ on murine splenic activated CD8^+^ T cells.

n=6 per group. Data are represented as mean ± SEM except for panel G. Unpaired two-tailed Student’s t-tests were used except for panels A, D, and G. Multiple unpaired t-test corrected by the Holm-Šidák method was used for panels A and D. Statistical significance was defined as follows: *p<0.05, **p<0.01, ***p<0.001, and ****p<0.0001.

**Figure S7. Cyst(e)ine supplementation stimulates lipid peroxidation and ROS levels in T-cells, related to Figure 4**

A. Representative T-distributed Stochastic Neighbor Embedding (t-SNE) plots on the Liperfluo channel of 6-day-activated T cells.

(B-G) Mean fluorescence intensity (MFI) of Liperfluo levels in murine splenic activated CD4^+^ (B-D) or CD8^+^ (E-G) T cells.

Representative t-SNE plots on the CellRox Deep Red channel of 6-day-activated T cells.
(I-N) MFI of ROS levels in murine splenic activated CD4^+^+ (I-K) or CD8^+^ (L-N) T cells.

n=6 per group, except for panels D, G, K, and N, n=5. Data are represented as mean ± SEM, whereas ***p<0.001 and ****p<0.0001 by unpaired two-tailed Student’s t-tests (B-G and I-N).

## STAR⍰METHODS

### EXPERIMENTAL MODEL AND STUDY PARTICIPANT DETAILS

#### Animal studies

The mice used in the PDAC models, including the orthotopic allograft murine KPC models and the KC model, and the C57BL/6J wild type mice (Jackson Laboratory, 000664) used for murine splenic T cell isolation were approved by the Institutional Animal Care and Use Committee (IACUC) at the University of Virginia and New York University, respectively. All mice were housed under specific pathogen-free (SPF) conditions under a 12-hour light-dark cycle and were randomly assigned to each experimental group.

For the orthotopic allograft murine PDAC models, both genders of C57BL/6J wildtype mice were purchased from the Jackson Laboratory (000664) and then bred on our own in the vivarium at the University of Virginia and New York University.

For the KC mouse model, C57BL/6J background transgenic p48-Cre mice and LSL-Kras^G12D/+^ mice were kindly gifted by Dr. Bar-Sagi Dafna (New York University Langone Health, NY). Genomic DNA of the offspring was isolated using the PCRBIO Rapid Extract Kit (PCR Biosystems, PB10.24-40). Genotyping was performed by polymerase chain reaction (PCR) on a thermal cycler (Eppendorf, Mastercycler X40) using the following primer pairs: p48-Cre (forward: 5’-ATAGGCTACCTGGCCATGCCC-3’, reverse: 5’-CGGGCTGCAGGAATTCGTCG-3’), LSL-Kras^G12D/+^ (forward: 5’-AGCTAGCCACCATGGCTTGAGTAAGT-3’, reverse: 5’-CCTTTACAAGCGCACGCAGAC-3’). PCR amplification was carried out with an annealing temperature of 60°C and an extension temperature of 72°C. The PCR products were separated by 2% agarose gel (Sigma-Aldrich, A9539) electrophoresis and visualized by iBright Imaging Systems (Thermo Fisher Scientific).

#### Orthotopic allograft murine PDAC model

KPC cells were harvested after trypsinization, thoroughly washed with pre-cold phosphate-buffered saline (PBS) three times, filtered through a 40 μm cell strainer, re-suspended in pre-cold PBS, and diluted 1:1 in pre-thawed phenol red-free matrigel (Corning, 356231) at a final concentration of 1 million/mL. The plungers of the 0.5 mL Lo-Dose™ Micro-Fine™ Insulin Syringes (Embecta, 329461) were pulled out, and 40 µL cell-matrigel suspension was loaded from the bottom of the syringes. The plungers were then reinserted into the syringes and pushed in to expel the air, until the cell-matrigel suspension was reserved at the 30 µL marking to control the injection volume, given 30K/mouse cancer cell implantation.

For the surgery of orthotopically implanting KPC cells, a pre-operative transdermal application of 40 mg/kg extended-release buprenorphine (Zorbium) will be administered 1 hour before surgery to ensure a less painful recovery from anesthesia. Mice were then anesthetized using isoflurane inhalation. The skin near the left upper quadrant of the abdomen was exposed by an electronic razor and pre-operatively prepped with three times 7.5% Povidone-Iodine scrub, and 70% alcohol. The pancreas and spleen were gently exposed through a 1-centimeter left transverse subcostal abdominal incision, and the 30 µL of the cell-matrigel suspension was slowly injected using the insulin syringe directly into the pancreas splenic lobe, resulting in a bright, soft, well-defined, non-leaking bleb. The pancreas and spleen were gently repositioned inside the abdomen, and the peritoneum was closed using 4-0 absorbable suture (Ethicon Vicryl, J845G). The skin was closed using stainless steel wound clips. Another dosage of Zorbium (40 mg/kg) was provided after 96 hours post-operative, and the clips were removed 14 days post-operative. Mice were monitored twice a week and euthanized at the pre-defined experimental endpoints or when they reached the humane welfare criteria, including exceeding 15% body weight loss, impaired mobility, abdominal distension, signs of pain or distress.

Mice at the endpoint were cardiac punctured to collect blood samples into microtainer blood collection tubes (BD Biosciences, 365967) under isoflurane anesthesia (5% induction and 1-2% maintenance) and then euthanized using inhaled isoflurane. The blood was centrifuged at 4,000 rpm for 15 minutes at 4°C, and the supernatant serum was collected and stored in a -80°C freezer. Tumors were harvested and weighed directly and stored in cold PBS for the following tumor-infiltrated lymphocytes extraction or in 4% paraformaldehyde (PFA) for fixation.

Rat IgG2a isotype control (5 mg/kg, Bioxcell, BP0089) or anti-mouse PD-1 (5 mg/kg, Bioxcell, BP0146) were intraperitoneally injected twice weekly in both the KPC and KC PDAC models.

*Akkermansia muciniphila* alone (2×10^9^/mL/kg) or combined with *Lacticaseibacillus reuteri* at 2 to 1 ratio (2×10^9^/mL/kg) dissolved in ddH_2_O was orally gavaged three times a week for the first two weeks, followed once a week for an additional four weeks for the combination groups of KPC and KC mice.

For the microbiome depletion experiment related to supplementary Figure S2, mice received a cocktail of vancomycin hydrochloride (50 mg/kg, Sigma-Aldrich, V2002), neomycin sulfate (100 mg/kg, Gibco, 21810031), and metronidazole (100 mg/kg, Sigma-Aldrich, PHR1052) by oral gavage three times a week for the first week, followed by ad libitum drinking water containing ampicillin sodium salt (1 mg/mL, Thermo Fisher Scientific, BP1760-25), vancomycin (0.5 mg/mL), neomycin (0.5 mg/mL) and metronidazole (1 mg/mL) for one week. The antibody injections and probiotic oral gavage were remaining same as previously described.

For the N-Acetyl-L-cysteine (NAC, Sigma-Aldrich, A9165) gavage procedure related to Figure 7, 50 mg/kg NAC was orally gavage twice a week for three or four weeks.

#### Cell lines and primary cultures

All cells were cultured in a 37°C incubator (Eppendorf, CellXpert ® C170i) maintaining 5% CO_2_. The KPC cell line was kindly gifted from Dr. Bar-Sagi Dafna from New York University Langone Health and cultured in Dulbecco’s Modified Eagle’s Medium (DMEM; Corning, 15-017-CV) supplemented with 10% heat-inactivated fetal bovine serum (FBS; Corning, 35-072-CV), 1× antibiotic antimycotic solution (Corning, 30-004-CI), and 2 mM L-alanyl-L-glutamine (Corning, 25-015-CI). The HEK293 cell line was gifted from Dr. Marco Davila (Roswell Park Comprehensive Cancer Center, Buffalo, NY) and cultured in DMEM/F-12 medium (Thermo Fisher Scientific, 11320033) supplemented with 10% FBS, 1% Penicillin-Streptomycin (Thermo Fisher Scientific, 15140122).

## METHOD DETAILS

### A. muciniphila and L. reuteri production

*A. muciniphila (Akkermansia muciniphila*; ATCC, BAA-835) was cultured under strict anaerobic conditions in brain heart infusion broth (Fisher Scientific, B99070) at 37°C for 24-48 hours. *L. reuteri* (*Lactobacillus reuteri*; ATCC, 23272) was revived from -80°C glycerol stocks and cultured in Mann-Rogosa-Sharpe (MRS) broth (Fisher Scientific, DF0881175) at 37°C under anaerobic conditions for 18-24 hours. Log-phase cultures (an OD600 value between 0.6-0.8) were collected anaerobically and washed in sterile pre-reduced PBS^45–47^. Formulation was prepared to enhance gastric survival and intestinal release. *A. muciniphila* and *L. reuteri* are microencapsulated within an alginate-gellan matrix prior to incorporation into xanthan gel. Briefly, bacterial suspensions are mixed with 1.5% sodium alginate (MP Biomedicals, ICN218295) and extruded into 100 mM calcium chloride (MP Biomedicals, ICN19381950) to form beads, followed by coating with 0.2% chitosan (Spectrum Chemical Manufacturing Corporation, 18-601-571) for acid resistance. Beads are incorporated into a practical grade xanthan (0.3%)-gellan (0.15%) hydrogel (MP Biomedicals, ICN96002180) containing 3% D-trehalose (Thermo Fisher Scientific, 309870250) and 2% inulin (MP Biomedicals, ICN19897180). Final pH is adjusted to 6.0. Enteric protection ensures minimal release at pH 1.5-3.0 and targeted release at intestinal pH≥6.5. Formulations are packaged under nitrogen and stored at 4°C^48–51^. Viability was assessed by serial dilution and plating on mucin-supplemented anaerobic agar and MRS agar for *A. muciniphila* and *L. reuteri*, respectively. Colony-forming units (CFU) were quantified before and during storage at 4°C. Encapsulation efficiency and stability were calculated as the percentage of recoverable CFU relative to input counts.

### Shotgun metagenomic sequencing

Fecal samples were collected using sterile 1.5 ml Eppendorf tubes, flash frozen, and stored at -80 °C until further processing. Aliquots of homogenized samples were subjected to DNA extraction using a Qiagen QIAamp PowerFecal Pro DNA kit (Qiagen, 51804) according to the manufacturer’s instructions. Extracted DNA was quantified using Quant-iT PicoGreen dsDNA Assay Kit (Invitrogen, P7589) and checked for purity by NanoDrop spectrophotometer (Thermo Fisher Scientific). Shotgun metagenomic DNA libraries were constructed with a target of 100 ng of DNA per sample using Illumina DNA Prep containing the Illumina Purification Beads and IDT for Illumina DNA/RNA UD Indexes according to the manufacturer’s instructions, using 5 PCR cycles (Illumina, Doc#1000000025416 v10). Libraries were pooled in equimolar amounts, a 1.0% PhiX Control spike-in was applied, and the pooled library was sequenced on an Illumina NextSeq 1000 platform with a 300-cycle (2×150) P2 reagent kit (V3). Negative controls were handled as samples and consisted of water and buffer used in DNA extractions and dilutions. ZymoBIOMICS Fecal Reference with TruMatrix Technology (ZYMO Research, D6323) was included as a sequencing quality control to facilitate inter-study comparisons.

### Bioinformatics and statistical analysis

Paired-end raw sequencing reads were quality-controlled using KneadData (https://huttenhower.sph.harvard.edu/kneaddata/) to remove low-quality and host contamination reads against mouse (C57BL) reference genome. The resulting clean reads were then taxonomically profiled with MetaPhlAn 4.2.2^52^ using the mpa_vJan25_CHOCOPhlAnSGB_202503 database. HUMAnN 3.9^53^ was used to profile functional composition. Functional pathway abundances were summarized at the MetaCyc pathway level and normalized to counts per million (CPM) for downstream analyses. Within-subject delta change (Endpoint-Baseline) was calculated for each matched pair and used for downstream cross-group pathway-level comparisons.

Taxonomic and functional profiles, together with sample metadata, were imported into phyloseq^54^ objects (version 1.54), in R (4.5.2) to perform downstream analysis. Observed and Shannon Alpha diversity metrics were calculated using the microbiome package (version 1.32), and differences between baseline and endpoint within each treatment group were assessed using a paired Wilcoxon signed-rank test. Beta diversity was assessed using Principal Coordinate Analysis (PCoA) based on Weighted UniFrac distance, Bray-Curtis dissimilarity, and Aitchison distance calculated from centered log-ratio (CLR) transformed relative abundances. For calculating Weighted UniFrac distance, CHOCOPhlAnSGB Newick tree is distributed with the same MetaPhlAn database used for profiling (mpa_vJan25_CHOCOPhlAnSGB_202503.nwk). Statistical differences in overall community composition were evaluated using permutational multivariate analysis of variance (PERMANOVA) via the adonis2^55,56^ function in the vegan package (version 2.7-2). Differentially abundant taxonomic and functional features were identified using the Wilcoxon rank-sum test. Species-species Spearman correlation analysis was performed using the Hmisc (version 5.2-5) package in R. Correlation heatmaps and network visualizations were generated using the pheatmap (version 1.0.13) and igraph (version 2.2.2) r packages. Differences were considered statistically significant at p<0.05.

### Tumor-infiltrated lymphocyte extraction

Tumor tissues were weighed by a balance scale (Denver Instrument, XL-3100) and minced into small segments using sterile iris scissors in gentleMACS™ C tubes (Miltenyi Biotec,130-096-334) containing 6-fold volumes of enzyme mix prepared according to the manufacturer’s instructions using the mouse Tumor Dissociation Kit (Miltenyi Biotec, 130-096-730). The C tubes were loaded onto a GentleMACS Octo Dissociator (Miltenyi Biotec, 130-134-029) and processed using the program 37C_m_TDK_2. The cell suspension after dissociation was smashed using the bottom of a 1 mL plunger, filtered into a 50 mL tube through a 100 µM cell strainer (Corning, CLS352360) using 10-fold volumes of PBS, and pelleted by centrifugation at 600 ×g for 4 minutes. The cell pellet was resuspended in 2 mL PBS, filtered again through a 100 µM cell strainer, and layered onto a Percoll (Sigma-Aldrich, GE17-0891-09) density gradient medium consisting of 80% and 40% Percoll solutions for the bottom and middle layers, respectively. After centrifuging at 2,000 rpm for 20 minutes at room temperature with minimal acceleration and braking, the third layer was carefully collected and washed twice with 12 mL PBS. The cell pellet was resuspended and cultured in 100 µL RPMI 1640 medium (1640; Corning, 15-040-CV) supplemented with 10% heat-inactivated FBS, 1× antibiotic antimycotic solution, 2 mM L-alanyl-L-glutamine, and 1× Brefeldin A solution (Biolegend, 420601) on a round-bottom 96-well plate for 6 hours.

### Mesenteric lymphocyte isolation

The entire mesenteric lymph node chain near the cecum, aligned with the colon, was exposed and harvested in sterile PBS. The lymphocytes were gently dissociated through a 100 µM cell strainer using the plunger of a 1 mL syringe and collected by centrifugation at 600 ×g for 4 minutes. The cells were resuspended in 1 mL PBS, and 50 µL cell suspension was plated for flow cytometry analysis.

### Flow cytometry analysis

Cell suspensions, including tumor-infiltrated lymphocytes, mesenteric lymphocytes, murine splenic T cells, human peripheral blood mononuclear cells (PBMC)-isolated T cells, and GFP CD19-28ζ CAR T cells were washed once with PBS and stained with Zombie UV™ dye (1:1000; Biolegend, 423108) for 15 minutes at room temperature in the dark. Then the cells were PBS washed, blocked in the TruStain FcX™ (anti-mouse CD16/32) Antibody (1:100; Biolegend, 101320) or the Human TruStain FcX™ (Fc Receptor Blocking Solution) (1:100; Biolegend, 422301) for murine samples and human samples, respectively, for 10 minutes at room temperature, and stained with a cocktail of surface markers (1:800 dilutions of Brilliant Violet 650™ anti-mouse CD45 Antibody (Biolegend, 103151), Brilliant Ultra Violet™ 496 CD3 Monoclonal Antibody (Thermo Fisher Scientific, 364-0032-82), Brilliant Violet 750™ anti-mouse CD4 Antibody (Biolegend, 100467), Spark Blue™ 550 anti-mouse CD8a Antibody (Biolegend, 100779), Brilliant Violet 785™ anti-mouse CD25 Antibody (Biolegend, 102051), Spark NIR™ 685 anti-mouse CD69 Antibody (Biolegend, 104558), PE/Dazzle™ 594 anti-mouse/human CD44 Antibody (Biolegend, 103056), Brilliant Violet 510™ anti-mouse CD62L Antibody (Biolegend, 104441), and BUV395 Rat Anti-Mouse CD279 (PD-1) Antibody (BD Biosciences, 568596) for murine samples, 1:400 dilutions of Brilliant Ultra Violet™ 805 CD4 Monoclonal Antibody (Thermo Fisher Scientific, 368-0049-42), Brilliant Violet 785™ anti-human CD8 Antibody (Biolegend, 344739), and APC/Cyanine7 anti-human CD279 (PD-1) Antibody (Biolegend, 329921) for human samples) for 20 minutes at room temperature in the dark. After twice PBS wash, cells were fixed and permeabilized using Cyto-Fast™ Fix/Perm Solution (Biolegend, 426803) for 20 minutes at room temperature in the dark. The cells were washed twice with 1× Perm Wash Solution and stained with an intracellular antibody cocktail (1:800 dilutions of Alexa Fluor® 700 anti-mouse TNF-α Antibody (Biolegend, 506338) and PerCP/Cyanine5.5 anti-mouse IFN-γ Antibody (Biolegend, 505822) for murine samples, while of Brilliant Violet 650™ anti-human TNF-α Antibody (Biolegend, 502937) and PE anti-human IFN-γ Antibody (Biolegend, 506506) for human samples) for another 20 minutes at room temperature in the dark. After twice PBS washing, the cells were resuspended in 1% PFA and analyzed on the 5-laser full spectral flow cytometer Cytek® Aurora (Cytek) within 3 days. The percentage of focused subsets and the mean fluorescence intensity (MFI) were quantified by FlowJo^TM^ 10.10.0 (BD Biosciences). The t-Distributed Stochastic Neighbor Embedding (t-SNE) platform in FlowJo^TM^ 10.10.0 was used for the t-SNE calculations. Cell numbers were normalized and calculated based on the tumor weight used in the flow cytometry.

### Immunohistochemistry and immunofluorescence

The KPC tumor blocks were fixed in 10% buffered formalin (Mercedes Scientific, MER44991GL) for 24 hours and then washed three times in 1× PBS, followed by automatic tissue processing (Leica, HistoCore PELORIS 3 Premium Tissue Processing System) for paraffin embedding. The paraffin blocks were sectioned at 5 μm thickness using a Leica rotary microtome.

For the immunofluorescence staining of CD4 and Ly-6G, KPC tumor sections were stained on a Leica BondRx auto-stainer according to the manufacturer’s instructions. In brief, sections were deparaffinized, followed by antigen retrieval with BOND Epitope Retrieval Solution 1 (pH6; Leica, AR9961) retrieval buffer at 100°C for 60 minutes for Ly-6G staining or BOND Epitope Retrieval Solution 2 (pH9; Leica, AR9640) for 20 minutes for CD4 staining. Sections were then treated with 3% Hydrogen Peroxide (RICCA, R3821310-1BV) to inhibit endogenous peroxidases. Slides were blocked with Immunofluorescence Blocking Buffer (Cell Signaling Technology, 12411) Blocker for CD4 and Rodent Block M (Biocare Medical, RBM961) for Ly-6G. After blocking, slides were incubated with the CD4 (D7D2Z) Rabbit Monoclonal Antibody (1:200; Cell Signaling Technology, 25229S) and Purified Rat Anti-Mouse Ly-6G (1:250; BD Biosciences, 551459) for 30 minutes and 60 minutes, respectively. Then the slides were incubated with Rabbit-on-Rodent HRP polymer (Biocare Medical, RMR622L) for CD4 and Rat-1-step HRP polymer (Biocare Medical, GHP516) for Ly-6G for 10 minutes. Following secondary incubation, slides underwent HRP-mediated tyramide signal amplification with Opal® 520 Reagent Pack (1:150; Akoya Biosciences, FP1487001KT) and Opal® 620 Reagent Pack (1:150; Akoya Biosciences, FP1495001KT) for CD4 and Ly-6G, respectively, for 10 minutes. Once the Opal® fluorophores were covalently linked to the antigen, primary and secondary antibodies were removed with a 95°C retrieval step. This sequence was repeated two more times with subsequent primary and secondary antibody pairs, using a different Opal® fluorophore with each primary antibody. After antibody staining, sections were counterstained with spectral DAPI (Akoya Biosciences, FP1490) and mounted with ProLong™ Gold Antifade Mountant (Thermo Fisher Scientific, P36935). Semi-automated image acquisition was performed on an Akoya Vectra Polaris (PhenoImagerHT) multispectral imaging system. Slides were scanned at 20X magnification using PhenoImagerHT 2.0.

For the immunohistochemistry staining of CD8, sections were stained on a Leica BondRX automated stainer, according to the manufacturer’s instructions. In brief, tissues underwent deparaffinization online followed by epitope retrieval for 20 minutes at 100°C with BOND Epitope Retrieval Solution 2 (pH9) and endogenous peroxidase activity blocking with 3% Hydrogen Peroxide. Sections were then incubated with CD8 alpha (D4W2Z) Rabbit Monoclonal Antibody (Cell Signaling Technology, 98941) at a 1:400 dilution for 30 minutes at room temperature. Primary antibodies were detected with anti-rabbit HRP-conjugated polymer and 3,3’-diaminobenzidine (DAB) substrate, both provided in the Leica BOND Polymer Refine Detection System (Leica, DS9800). Finally, samples were counterstained with hematoxylin (Biocare, CATHE-M), dehydrated, and coverslipped with Permount™ Mounting Medium (Electron Microscopy Sciences, 17986). Slides were scanned at 40X on a Hamamatsu Nanozoomer (2.0HT) whole slide scanner, and the image files were uploaded to the NYUGSoM’s OMERO Plus image data management system (Glencoe Software).

The data was quantified by HALO (Indica Labs), an AI-powered, digital pathology image analysis platform. Briefly, the platform was trained to identify every single cell and stained-positive cell, thus allowing it to export the numbers of total cells and stained-positive cells from the whole slide.

### Cysteine level measurement

Cysteine levels in serum and tumor tissue were measured using the Cysteine Assay Colorimetric Kit (Novus Biologicals, NBP3-25785). Serum was diluted 10 times using reagent 1 provided by the kits, and centrifuged at 10000 ×g for 10 min at 4°C. Tumor tissue samples were combined with 9-fold volume of reagent 1, mechanically lysed using TissueLyser III (QIAGEN, 9003240), then centrifuged at 10000 ×g for 10 min at 4°C. The reactions containing 20 µL of the samples from supernatant, 100 µL of reagent 2, and 100 µL of reagent 3 were performed in a 96-well microplate at room temperature for 10 minutes. The absorbance at 600 nm was measured with a microplate reader (Thermo Fisher Scientific, Varioskan LUX).

### Isolation of murine splenic T cells

Spleens from 8-week-old C57BL/6J wild type male mice were harvested in PBS, smashed using the bottom of 1 mL plungers on 100 µM strainers, flushed with PBS through the strainer, and spun down at 600 ×g for 4 min at 4°C. The cell pellet was treated with 500 µL 1× red blood cell lysis buffer (Biolegend, 420302) for 2 minutes at room temperature, filled with 5 mL PBS, and spun down at 600 ×g for 4 min at 4°C. The cells were then resuspended with 1 mL PBS, filtered through 100 µM strainers, and T cell isolation was performed using the EasySep™ mouse T cell Isolation Kit (Stemcell, 19851). Basically, 1 mL of splenocytes (less than 10^8^) was incubated with 20 µL FcR blocker and 50 µL of isolation cocktail for 10 minutes at room temperature in flow tubes (Thomas Scientific, 23A00A610) to have unwanted cells labeled with biotinylated antibodies, and then 75 µL RapidSpheres™ was added and incubated for 2.5 minutes at room temperature to bind the labeled unwanted cells. Then, flow tubes were topped with 1.5 mL complete 1640 medium, gently pipetting up and down several times, and placed into the magnet for 2.5 minutes at room temperature. The supernatant containing purified T cells was collected, counted by TC20 automated cell counter (Bio-Rad), and plated at a density of 0.1 million/100 µL/well in a round-bottom 96-well plate for unactivation cultures, or in an anti-CD3-coated flat-bottom 96-well plate for activation cultures.

### Human T cell isolation

PBMCs were isolated from a healthy donor’s whole blood using lymphocyte separation medium (Corning, 25-072-CI) and collected from the lymphocyte layer. Cells were washed, resuspended in a PBMC dilute solution (Dulbecco’s PBS (DPBS; Thermo Fisher Scientific, 14040-133), 2% FBS, and 1mM EDTA (Thermo Fisher Scientific, AM9260G)), counted, and used immediately or stored in a frozen medium consisting of 90% FBS and 10% DMSO.

Human T cells were isolated from PBMCs using the EasySep™ Human T Cell Isolation Kit (Stemcell, 17951) according to the manufacturer’s instructions. Purified T cells were cultured in high-glucose RPMI 1640 medium (Thermo Fisher Scientific, A1049101) supplemented with 10% FBS, 1% Penicillin-Streptomycin, and 200 U/mL human IL-2 recombinant protein (Thermo Fisher Scientific, 200-02-1MG) (defined as human T cell medium below), and activated using human T-Activator CD3/CD28 magnetic beads (Thermo Fisher Scientific, 11131D) at a 1:1 beads to cells ratio for 48 hours.

### Generation of GFP CD19-28**ζ** CAR T cells

Retroviral supernatant was harvested from the medium of HEK293 cells that produced supernatant of the virus loaded with GFP CD19-28ζ CAR vectors. Non-tissue culture-treated 6-well plates were coated with RetroNectin reagent (Takara, T100A) diluted in DPBS (15 µg/ml) overnight at 4°C, blocked with 0.5% DPBS-disolved BSA, and washed before use. 48-hour-activated human PBMC-isolated T cells were plated at a density of 3-5×10^6^ per well and incubated with retroviral supernatant in human T cell medium in the 6-well RetroNectin-coated plate and centrifuged at 2,000 ×g for 1 hour at room temperature. T cells were incubated in the 37°C incubator for 24 hours and followed by a second transduction the following day. Over 90% positive transduction efficiency was verified by flow cytometry and quantified based on the expression of GFP. Transduced CAR T-GFP cells were subsequently expanded in human T cell medium for flow cytometry assays.

### Murine and human T cell activation assay

For murine splenic T cell activation experiments, 100 μL 2 μg/mL Purified anti-mouse CD3 Antibody (Biolegend, 100238) was coated on the flat-bottom 96-well plate in the 37°C incubator for 2 hours. Isolated murine T cells were plated at a density of 0.1 million/100 µL/well and activated in cyst(e)ine-free 1640 medium (Sigma-Aldrich, R7513-100ML) containing 10% FBS, 1× antibiotic antimycotic solution, 2 mM L-alanyl-L-glutamine, 100 μM methionine (Sigma-Aldrich, M9625), and 5 μg/mL Purified anti-mouse CD28 Antibody (Biolegend, 102121). T cell numbers and viabilities were measured by the TC20 automated cell counter based on trypan blue staining (Thermo Fisher Scientific, 15250061), and T cell activation levels were evaluated by flow cytometry at the indicated timepoints.

For human T cell activation experiments, isolated human T cells or GFP CD19-28ζ CAR T cells were recovered from the nitrogen stocks and fully rested by culture in human T cell medium after removing the CD3/CD28 magnetic beads for 14 days. 0.3 million/mL/well human T cells in a 24-well flat-bottom plate were plated and re-activated by adding human T-Activator CD3/CD28 magnetic beads at a 1:1 bead-to-cell ratio for 48 hours in cyst(e)ine-free RPMI 1640 medium supplemented with 10% FBS, 1% Penicillin-Streptomycin, 2 mM L-alanyl-L-glutamine, 100 μM methionine, 10mM HEPES (Goldbio, H-400-100) and 2.5 g/L D-glucose (Sigma-Aldrich, G8270) for the following culturing or for flow cytometry analysis.

### Cyst(e)ine additives

20 μM L-Cystine dihydrochloride (CySS, Sigma-Aldrich, C6727) dissolved in 2 M HCl, supplied with PBS or 400 μM N-Acetyl-L-cysteine (Cys, Sigma-Aldrich, A9165), was used for Figures 4-6 and Figures S3C and S4. 200 μM CySS dissolved in 2 M HCl (HCl as the control) was used to administer CySS levels in Figures S3E and S5, while 400 μM Cys dissolved in PBS (PBS as the control) was used to administer Cys levels in Figures S3D and S6. Conditions in Figure S7 were administered as mentioned.

### RNA isolation, reverse-transcription, and RT-qPCR

Total RNA was extracted from murine splenic T cells using the RNeasy Plus Mini Kit (QIAGEN, 74136), following the manufacturer’s protocol. The concentration and the quality of the RNA were evaluated by the NanoDrop spectrophotometer (Thermo Fisher Scientific). 1 μg of RNA was reverse transcribed using the iScript cDNA Synthesis Kit (Bio-Rad, 1708891) in a total volume of 20 μL reaction mixture. RT-qPCR (real-time quantitative PCR) was performed on a QuantStudio™ 5 system (Thermo Fisher Scientific) using PowerTrack™ SYBR Green Master Mix (Thermo Fisher Scientific, A46109). The reactions were carried out in a total volume of 10 μL mixture containing cDNA, 100 nmol/L of both forward and reverse primers, and Power SYBR Green Master Mix in 384-well clear reaction plates (Bio-Rad, A36931). The cycling conditions were 94°C for 10 min, followed by 40 cycles of 94°C for 30 seconds and 60°C for 30 seconds, and a final elongation at 72°C for 5 min. Actb served as the housekeeping gene. The 2^-ΔΔCT^ method was used for data analysis, and the results were presented as relative mRNA expression levels. The RT-qPCR primers used in this study are listed below:

Actb Forward (F): 5’-GGCTGTATTCCCCTCCATCG-3’, Reverse (R): 5’-GCACAGGGTGCTCCTCAG-3’; Ki67 F: 5’-GAGGAGAAACGCCAACCAAGAG-3’, R: 5’-TTTGTCCTCGGTGGCGTTATCC-3’; Ccna2 F: 5’-TTGTAGGCACGGCTGCTATGCT-3’, R: 5’-GGTGCTCCATTCTCAGAACCTG-3’; Ccnb1 F: 5’-AGAGGTGGAACTTGCTGAGCCT-3’, R: 5’-GCACATCCAGATGTTTCCATCGG-3’; Ccne2 F: 5’-GTACTGTCTGGAGGAATCAGCC-3’, R: 5’-CCAAACCTCCTGTGAACATGCC-3’; Slc1a1 F: 5’-GGAAGAACCCTTTCCGCTTTGC-3’, R: 5’-CACAGCGGAATGTAACTGGCAG-3’; Slc1a4 F: 5’-GAGGGAGAAGACCTCATCCGAT-3’, R: 5’-GTCACCAGCATGACGATGTCCT-3’; Slc1a5 F: 5’-CTGCCTGTGAAGGACATCTCCT-3’, R: 5’-CTCGGCATCTTGGTTCGATCCA-3’; Slc3a2 F: 5’-GAGCGTACTGAATCCCTAGTCAC-3’, R: 5’-GCTGGTAGAGTCGGAGAAGATG-3’; Slc7a9 F: 5’-GGATTCCTCTGGTGACCGTATG-3’, R: 5’-CAAGATGCTGGATAGAGAACGCG-3’; Slc7a10 F: 5’-TGCTACGGAGTCACTATCCTGG-3’, R: 5’-GCTGAAGACCAGTAGGAATGCC-3’; Slc7a11 F: 5’-AGGGCATACTCCAGAACACG-3’, R: 5’-GGACCAAAGACCTCCAGAATG-3’.

### Half-maximal effective concentration (EC_50_) assay

Cell counting kit-8 (CCK-8; ApexBio Technology, K1018) assay was used to determine the half maximal effective concentration of CySS/Cys on T cell or KPC cell growth. For T cells, isolated cells were plated at a density of 0.1 million/100 μL/well and cultured in cyst(e)ine-free 1640 medium supplemented with 10% FBS, 1× antibiotic antimycotic solution, 2 mM L-alanyl-L-glutamine, 100 μM methionine, and graded concentrations of CySS/Cys, with or without anti-CD3 plus anti-CD28, to generate activated (in flat bottom 96-well plates) or unactivated (in round bottom 96-well plates) T cells, respectively. For KPC cells, 15000/well cells were seeded onto flat-bottom 96-well plates and cultured in complete DMEM medium overnight. The medium was replaced the following day with 100 μL cyst(e)ine-free DMEM medium (Sigma-Aldrich, D0422-100ML) supplemented with 10% FBS, 1× antibiotic antimycotic solution, 2 mM L-alanyl-L-glutamine, 100 μM methionine, and graded concentrations of CySS/Cys. 10 μL CCK-8 reagents were added in each well at 48 hours (for unactivated T cells) or 72 hours (for activated T cells and KPC cells) and incubated at 37°C for 1 hour. The absorbance at 450 nm was recorded using the Varioskan LUX microplate reader (Thermo Fisher Scientific).

### KPC cell killing and crystal violet staining

Isolated T cells were activated in cyst(e)ine-free 1640 medium supplemented with 10% FBS, 1× antibiotic antimycotic solution, 2 mM L-alanyl-L-glutamine, 100 μM methionine, 5 μg/mL anti-CD28, and 20 μM of CySS, as the control, or 20 μM of CySS plus 400 μM of Cys on day 1 for 3 days. On day 3, 2,000/well KPC cells were seeded onto flat-bottom 96-well plates and cultured in complete DMEM medium overnight. On day 4, the KPC cells from representative wells were counted. Meanwhile, T cells from each group were harvested, counted, PBS-washed once, and resuspended in complete 1640 medium. KPC cell medium was replaced with 100 μL T cell suspension at a 2:1 effector to target ratio.

After co-culturing KPC cells and T cells for 48 hours, the KPC cells killed by T cells were evaluated by staining the adherent KPC cells with crystal violet. The medium was removed, and the T cells were gently washed out 3 times with PBS. The adherent cells were fixed with methanol in the fume hood and covered with 50% glycerol (Sigma-Aldrich, G5516) at room temperature overnight. After twice PBS washing, the wells were stained with 100 μL 0.05% crystal violet (Sigma-Aldrich, C0775) for 10 minutes at room temperature, followed by 5 times washes with PBS. The wells were completely dried and photographed using a microscope (Leica, Ivesta_3I 4.00a). QuPath v.0.6.0 (https://qupath.github.io/) was used to quantify the crystal violet-stained area^57^. In each well, regions with a threshold value greater than 60 (resolution=full, smoothing sigma=1) were classified as positive area. The total area and the stained-positive area for each well were quantified and exported for subsequent analysis. The parameters were applied to all images to ensure consistency and comparability.

### Lipid peroxides detection

Liperfluo (Cayman Chemical, 1448846-35-2) was used to detect lipid peroxides in T cells following the manufacturer’s instructions. Briefly, T cells were transformed into serum-free 1640 containing 1 μmol/L Liperfluo and incubated at 37°C for 30 minutes. The liperfluo fluorescence was detected by flow cytometry at 488 nm for the excitation wavelength and 515-545 nm for the emission wavelength.

### Cellular reactive oxygen species (ROS) measurement

CellROX™ Deep Red Reagent (Thermo Fisher Scientific, C10422) was used to measure the cellular ROS level. T cells were incubated with 5 μM CellROX™ reagent at 37°C for 30 minutes and then washed with PBS twice and fixed in 4% PFA. The CellROX™ Deep Red fluorescence was detected by flow cytometry at 640 nm for the excitation wavelength and 665 nm for the emission wavelength.

### Bulk RNA sequencing analysis

Isolated murine splenic T cells were activated for 3 days with 20 μM of CySS, supplemented with PBS or 400 μM of Cys. Total RNA was extracted as previously described, and the library preparation and the run of RNA-seq were performed by the University of Virginia Genome Analysis Technology Core using standard protocols. Briefly, libraries were prepared using NEBNext UltraExpress® RNA Library Prep Kit (NEW ENGLAND Biolabs, E3330L) and NEBNext® Poly(A) mRNA Magnetic Isolation Module (NEW ENGLAND Biolabs, E7490L). Libraries were sequenced on Illumina NextSeq 2000 Sequencing System (Illumina, 20038897) using NextSeq™ 1000/2000 P2 XLEAP-SBS™ Reagent Kit (100 Cycles) (Illumina, 20100987) in paired-end mode. FASTQ files were processed using the Salmon tool and the Tximport tool, and the differentially expressed genes between the groups were analyzed and exported using the DESeq2 tool available in the Galaxy platform^58^ (usegalaxy.org). Differentially expressed genes in Cys-supplement group were ranked by signed log2(fold change) multiplied by -log10(p-value) to generate the pre-ranked gene list. Gene set enrichment analysis (GSEA) was then performed using the M2:CP:REACTOME gene sets with the GESAPreranked tool in the GSEA software (v.4.4.0), based on the pre-ranked gene list^59,60^.

### EdU tracing, EU tracing, and puromycin tracing

T cells were incubated with 10 μM 5-ethynyl-2’-deoxyuridine (EdU) or 1 mM 5-ethynyl uridine (EU), provided from Click-iT™ EdU cell proliferation kit (Thermo Fisher Scientific, C10337) and Click-iT® RNA imaging kit (Thermo Fisher Scientific, C10329), respectively, or 10 μg/mL puromycin dihydrochloride (Sigma-Aldrich, A1113802) for 1 hour at 37°C. The cells were PBS-washed once, stained for viability using Zombie UV™ Fixable Viability Kit (1:1000, Biolegend, 423108) for 15 minutes at room temperature in the dark, and fixed and permeabilized using True-Nuclear™ transcription factor buffer set (Biolegend, 424401) for 1 hour at room temperature in the dark. EdU and EU labeled cells were incubated with freshly made Click-iT® reactions contained Click-iT® EdU buffer additive (1:10), CuSO_4_ (1:50), Alexa Fluor® azide (1:200), while puromycin dihydrochloride labeled cells were stained with Alexa Fluor® 488 anti-puromycin antibody (1:800, Biolegend, 381505), for 30 minutes at room temperature in the dark. After twice washing with PBS, the Alexa Fluor® 488 signal was detected by Cytek® Aurora, and the percentage of Alexa Fluor® 488-positive cells among alive cells was quantified by FlowJo^TM^ 10.10.0 (BD Biosciences).

## QUANTIFICATION AND STATISTICAL ANALYSIS

### Statistical analysis

All statistical analyses were performed using GraphPad Prism 10 software, and all values in the study are presented as the mean ± standard error of the mean (SEM). Unpaired two-tailed Student’s t-tests were used to compare two groups. One-way analysis of variance (ANOVA) followed by Fisher’s LSD tests and by Tukey’s post hoc tests were used for the comparisons among three and four groups, respectively. The Pearson correlation coefficient was used to assess the strength and direction of the relationship for Figures 3C and 3E. Multiple unpaired t-tests corrected by the Holm-Šidák method were used for growth curves (panels A and D in Figures 4, S5, and S6) and Figures 6F-6I. Statistical significance was defined as follows: ns p>0.05, *p<0.05, **p<0.01, ***p<0.001, and ****p<0.0001. The exact sample numbers (n for murine samples, N for human samples) and statistical analysis are indicated in each figure legend.

**Table S1. Spearman correlations between Akkermansia muciniphila and other species in all treatment naive baseline and combination group endpoint samples. Related to** Figure 1**. Spearman’s rank correlation coefficient (r) and corresponding p values are reported for baseline (Spearman_r_1, P_value_1) and endpoint (Spearman_r_2, P_value_2).**

**Table S2. Significant MetaCyc pathways identified by Wilcoxon testing (p<0.05), comparing the delta change in pathway CPM between the Anti-PD-1 and Combination groups. Related to** Figure 3.

**Table S3. Cyst(e)ine transporters in qPCR analysis. Related to Figure S3.**

**Table S4. Half-maximal growth concentration (EC_50_) of cyst(e)ine in unactivated T cells, activated T cells, and KPC cells. Related to Figure S3.**

**Table S5. Numbers and percentages of positively and negatively enriched pathways in each hierarchy following cysteine supplementation. Related to** Figure 5.

## Notes

https://data.mendeley.com/preview/wgzh4pzw5p?a=68683e00-9748-40ff-80fd-67be0a790806

https://data.mendeley.com/preview/7bn53gnsnk?a=6041a4c5-e097-4f35-8a57-8d6763c2cce6

### Competing Interest Statement

The authors have declared no competing interest.

## REFERENCES

1. Henriksen, A., Dyhl-Polk, A., Chen, I., and Nielsen, D. (2019). Checkpoint inhibitors in pancreatic cancer. Cancer Treat Rev 78, 17–30. 10.1016/j.ctrv.2019.06.005.

2. O’Reilly, E.M., Oh, D.Y., Dhani, N., Renouf, D.J., Lee, M.A., Sun, W., Fisher, G., Hezel, A., Chang, S.C., Vlahovic, G., et al. (2019). Durvalumab With or Without Tremelimumab for Patients With Metastatic Pancreatic Ductal Adenocarcinoma: A Phase 2 Randomized Clinical Trial. JAMA Oncol 5, 1431–1438. 10.1001/jamaoncol.2019.1588.

3. Ajina, R., Malchiodi, Z.X., Fitzgerald, A.A., Zuo, A., Wang, S., Moussa, M., Cooper, C.J., Shen, Y., Johnson, Q.R., Parks, J.M., et al. (2021). Antitumor T-cell Immunity Contributes to Pancreatic Cancer Immune Resistance. Cancer Immunol Res 9, 386–400. 10.1158/2326-6066.CIR-20-0272.

4. Lu, Y., Yuan, X., Wang, M., He, Z., Li, H., Wang, J., and Li, Q. (2022). Gut microbiota influence immunotherapy responses: mechanisms and therapeutic strategies. J Hematol Oncol 15, 47. 10.1186/s13045-022-01273-9.

5. Malczewski, A.B., Coward, J.I., Ketheesan, N., and Navarro, S. (2025). Immunometabolism: The role of gut-derived microbial metabolites in optimising immune response during checkpoint inhibitor therapy. Clin Transl Med 15, e70472. 10.1002/ctm2.70472.

6. Mimpen, I.L., Battaglia, T.W., Parra-Martinez, M., Toner-Bartelds, C., Zeverijn, L.J., Geurts, B.S., Verkerk, K., Hoes, L.R., van Renterghem, A.W.J., Noe, M., et al. (2026). Microbial Metabolic Pathways Guide Response to Immune Checkpoint Blockade Therapy. Cancer Discov 16, 95–113. 10.1158/2159-8290.CD-24-1669.

7. Peng, Z., Cheng, S., Kou, Y., Wang, Z., Jin, R., Hu, H., Zhang, X., Gong, J.F., Li, J., Lu, M., et al. (2020). The Gut Microbiome Is Associated with Clinical Response to Anti-PD-1/PD-L1 Immunotherapy in Gastrointestinal Cancer. Cancer Immunol Res 8, 1251–1261. 10.1158/2326-6066.CIR-19-1014.

8. Rasouli, B.S., Ghadimi-Darsajini, A., Nekouian, R., and Iragian, G.R. (2017). In vitro activity of probiotic Lactobacillus reuteri against gastric cancer progression by downregulation of urokinase plasminogen activator/urokinase plasminogen activator receptor gene expression. J Cancer Res Ther 13, 246–251. 10.4103/0973-1482.204897.

9. Lee, J.Y., Han, G.G., Choi, J., Jin, G.D., Kang, S.K., Chae, B.J., Kim, E.B., and Choi, Y.J. (2017). Pan-Genomic Approaches in Lactobacillus reuteri as a Porcine Probiotic: Investigation of Host Adaptation and Antipathogenic Activity. Microb Ecol 74, 709–721. 10.1007/s00248-017-0977-z.

10. Braathen, G., Ingildsen, V., Twetman, S., Ericson, D., and Jorgensen, M.R. (2017). Presence of Lactobacillus reuteri in saliva coincide with higher salivary IgA in young adults after intake of probiotic lozenges. Benef Microbes 8, 17–22. 10.3920/BM2016.0081.

11. Newsome, R.C., Gharaibeh, R.Z., Pierce, C.M., da Silva, W.V., Paul, S., Hogue, S.R., Yu, Q., Antonia, S., Conejo-Garcia, J.R., Robinson, L.A., and Jobin, C. (2022). Interaction of bacterial genera associated with therapeutic response to immune checkpoint PD-1 blockade in a United States cohort. Genome Med 14, 35. 10.1186/s13073-022-01037-7.

12. Davis, S.R., Stacpoole, P.W., Williamson, J., Kick, L.S., Quinlivan, E.P., Coats, B.S., Shane, B., Bailey, L.B., and Gregory, J.F., 3rd (2004). Tracer-derived total and folate-dependent homocysteine remethylation and synthesis rates in humans indicate that serine is the main one-carbon donor. Am J Physiol Endocrinol Metab 286, E272–279. 10.1152/ajpendo.00351.2003.

13. Stipanuk, M.H. (2004). Sulfur amino acid metabolism: pathways for production and removal of homocysteine and cysteine. Annu Rev Nutr 24, 539–577. 10.1146/annurev.nutr.24.012003.132418.

14. Wang, W., and Zou, W. (2020). Amino Acids and Their Transporters in T Cell Immunity and Cancer Therapy. Mol Cell 80, 384–395. 10.1016/j.molcel.2020.09.006.

15. Parker, J.L., Deme, J.C., Kolokouris, D., Kuteyi, G., Biggin, P.C., Lea, S.M., and Newstead, S. (2021). Molecular basis for redox control by the human cystine/glutamate antiporter system xc(). Nat Commun 12, 7147. 10.1038/s41467-021-27414-1.

16. Angelini, G., Gardella, S., Ardy, M., Ciriolo, M.R., Filomeni, G., Di Trapani, G., Clarke, F., Sitia, R., and Rubartelli, A. (2002). Antigen-presenting dendritic cells provide the reducing extracellular microenvironment required for T lymphocyte activation. Proc Natl Acad Sci U S A 99, 1491–1496. 10.1073/pnas.022630299.

17. Levring, T.B., Hansen, A.K., Nielsen, B.L., Kongsbak, M., von Essen, M.R., Woetmann, A., Odum, N., Bonefeld, C.M., and Geisler, C. (2012). Activated human CD4+ T cells express transporters for both cysteine and cystine. Sci Rep 2, 266. 10.1038/srep00266.

18. Srivastava, M.K., Sinha, P., Clements, V.K., Rodriguez, P., and Ostrand-Rosenberg, S. (2010). Myeloid-derived suppressor cells inhibit T-cell activation by depleting cystine and cysteine. Cancer Res 70, 68–77. 10.1158/0008-5472.CAN-09-2587.

19. Karlsson, H., Nava, S., Remberger, M., Hassan, Z., Hassan, M., and Ringden, O. (2011). N-acetyl-L-cysteine increases acute graft-versus-host disease and promotes T-cell-mediated immunity in vitro. Eur J Immunol 41, 1143–1153. 10.1002/eji.201040589.

20. Delneste, Y., Jeannin, P., Potier, L., Romero, P., and Bonnefoy, J.Y. (1997). N-acetyl-L-cysteine exhibits antitumoral activity by increasing tumor necrosis factor alpha-dependent T-cell cytotoxicity. Blood 90, 1124–1132.

21. Chi, F., Zhang, Q., Shay, J.E.S., Han, S., Ten Hoeve, J., Yuan, Y., Yang, Z., Shin, H., Block, S., Solanki, S., et al. (2025). Dietary cysteine enhances intestinal stemness via CD8(+) T cell-derived IL-22. Nature 647, 706–715. 10.1038/s41586-025-09589-5.

22. Green, J.L., Heard, K.J., Reynolds, K.M., and Albert, D. (2013). Oral and Intravenous Acetylcysteine for Treatment of Acetaminophen Toxicity: A Systematic Review and Meta-analysis. West J Emerg Med 14, 218–226. 10.5811/westjem.2012.4.6885.

23. Pushalkar, S., Hundeyin, M., Daley, D., Zambirinis, C.P., Kurz, E., Mishra, A., Mohan, N., Aykut, B., Usyk, M., Torres, L.E., et al. (2018). The Pancreatic Cancer Microbiome Promotes Oncogenesis by Induction of Innate and Adaptive Immune Suppression. Cancer Discov. 10.1158/2159-8290.CD-17-1134.

24. Dao, M.C., Everard, A., Aron-Wisnewsky, J., Sokolovska, N., Prifti, E., Verger, E.O., Kayser, B.D., Levenez, F., Chilloux, J., Hoyles, L., et al. (2016). Akkermansia muciniphila and improved metabolic health during a dietary intervention in obesity: relationship with gut microbiome richness and ecology. Gut 65, 426–436. 10.1136/gutjnl-2014-308778.

25. Fan, S., Jiang, Z., Zhang, Z., Xing, J., Wang, D., and Tang, D. (2023). Akkermansia muciniphila: a potential booster to improve the effectiveness of cancer immunotherapy. J Cancer Res Clin Oncol 149, 13477–13494. 10.1007/s00432-023-05199-8.

26. Zhu, Y., Chen, B., Zhang, X., Akbar, M.T., Wu, T., Zhang, Y., Zhi, L., and Shen, Q. (2024). Exploration of the Muribaculaceae Family in the Gut Microbiota: Diversity, Metabolism, and Function. Nutrients 16. 10.3390/nu16162660.

27. Lagkouvardos, I., Lesker, T.R., Hitch, T.C.A., Galvez, E.J.C., Smit, N., Neuhaus, K., Wang, J., Baines, J.F., Abt, B., Stecher, B., et al. (2019). Sequence and cultivation study of Muribaculaceae reveals novel species, host preference, and functional potential of this yet undescribed family. Microbiome 7, 28. 10.1186/s40168-019-0637-2.

28. Mukherjee, S., Lodha, T.D., and Madhuprakash, J. (2023). Comprehensive Genome Analysis of Cellulose and Xylan-Active CAZymes from the Genus Paenibacillus: Special Emphasis on the Novel Xylanolytic Paenibacillus sp. LS1. Microbiol Spectr 11, e0502822. 10.1128/spectrum.05028-22.

29. Hingorani, S.R., Petricoin, E.F., Maitra, A., Rajapakse, V., King, C., Jacobetz, M.A., Ross, S., Conrads, T.P., Veenstra, T.D., Hitt, B.A., et al. (2003). Preinvasive and invasive ductal pancreatic cancer and its early detection in the mouse. Cancer Cell 4, 437–450. 10.1016/s1535-6108(03)00309-x.

30. Pinato, D.J., Howlett, S., Ottaviani, D., Urus, H., Patel, A., Mineo, T., Brock, C., Power, D., Hatcher, O., Falconer, A., et al. (2019). Association of Prior Antibiotic Treatment With Survival and Response to Immune Checkpoint Inhibitor Therapy in Patients With Cancer. JAMA Oncol 5, 1774–1778. 10.1001/jamaoncol.2019.2785.

31. Zhang, M., Liu, J., and Xia, Q. (2023). Role of gut microbiome in cancer immunotherapy: from predictive biomarker to therapeutic target. Exp Hematol Oncol 12, 84. 10.1186/s40164-023-00442-x.

32. Hoentjen, F., Harmsen, H.J., Braat, H., Torrice, C.D., Mann, B.A., Sartor, R.B., and Dieleman, L.A. (2003). Antibiotics with a selective aerobic or anaerobic spectrum have different therapeutic activities in various regions of the colon in interleukin 10 gene deficient mice. Gut 52, 1721–1727.

33. Pushalkar, S., Hundeyin, M., Daley, D., Zambirinis, C.P., Kurz, E., Mishra, A., Mohan, N., Aykut, B., Usyk, M., Torres, L.E., et al. (2018). The Pancreatic Cancer Microbiome Promotes Oncogenesis by Induction of Innate and Adaptive Immune Suppression. Cancer Discov 8, 403–416. 10.1158/2159-8290.CD-17-1134.

34. McBean, G.J. (2002). Cerebral cystine uptake: a tale of two transporters. Trends Pharmacol Sci 23, 299–302. 10.1016/s0165-6147(02)02060-6.

35. Mosharov, E., Cranford, M.R., and Banerjee, R. (2000). The quantitatively important relationship between homocysteine metabolism and glutathione synthesis by the transsulfuration pathway and its regulation by redox changes. Biochemistry 39, 13005–13011. 10.1021/bi001088w.

36. Meister, A., and Anderson, M.E. (1983). Glutathione. Annu Rev Biochem 52, 711–760. 10.1146/annurev.bi.52.070183.003431.

37. Devadas, S., Zaritskaya, L., Rhee, S.G., Oberley, L., and Williams, M.S. (2002). Discrete generation of superoxide and hydrogen peroxide by T cell receptor stimulation: selective regulation of mitogen-activated protein kinase activation and fas ligand expression. J Exp Med 195, 59–70. 10.1084/jem.20010659.

38. Sena, L.A., Li, S., Jairaman, A., Prakriya, M., Ezponda, T., Hildeman, D.A., Wang, C.R., Schumacker, P.T., Licht, J.D., Perlman, H., et al. (2013). Mitochondria are required for antigen-specific T cell activation through reactive oxygen species signaling. Immunity 38, 225–236. 10.1016/j.immuni.2012.10.020.

39. Lin, J., Lee, I.M., Song, Y., Cook, N.R., Selhub, J., Manson, J.E., Buring, J.E., and Zhang, S.M. (2010). Plasma homocysteine and cysteine and risk of breast cancer in women. Cancer Res 70, 2397–2405. 10.1158/0008-5472.CAN-09-3648.

40. Zhang, S.M., Willett, W.C., Selhub, J., Manson, J.E., Colditz, G.A., and Hankinson, S.E. (2003). A prospective study of plasma total cysteine and risk of breast cancer. Cancer Epidemiol Biomarkers Prev 12, 1188–1193.

41. Hatae, R., Chamoto, K., Kim, Y.H., Sonomura, K., Taneishi, K., Kawaguchi, S., Yoshida, H., Ozasa, H., Sakamori, Y., Akrami, M., et al. (2020). Combination of host immune metabolic biomarkers for the PD-1 blockade cancer immunotherapy. JCI Insight 5. 10.1172/jci.insight.133501.

42. Lin, Z., Yang, S., Qiu, Q., Cui, G., Zhang, Y., Yao, M., Li, X., Chen, C., Gu, J., Wang, T., et al. (2024). Hypoxia-induced cysteine metabolism reprogramming is crucial for the tumorigenesis of colorectal cancer. Redox Biol 75, 103286. 10.1016/j.redox.2024.103286.

43. Cramer, S.L., Saha, A., Liu, J., Tadi, S., Tiziani, S., Yan, W., Triplett, K., Lamb, C., Alters, S.E., Rowlinson, S., et al. (2017). Systemic depletion of L-cyst(e)ine with cyst(e)inase increases reactive oxygen species and suppresses tumor growth. Nat Med 23, 120–127. 10.1038/nm.4232.

44. Gazzaniga, F.S., and Kasper, D.L. (2025). The gut microbiome and cancer response to immune checkpoint inhibitors. J Clin Invest 135 10.1172/JCI184321.

45. Plovier, H., Everard, A., Druart, C., Depommier, C., Van Hul, M., Geurts, L., Chilloux, J., Ottman, N., Duparc, T., Lichtenstein, L., et al. (2017). A purified membrane protein from Akkermansia muciniphila or the pasteurized bacterium improves metabolism in obese and diabetic mice. Nat Med 23, 107–113. 10.1038/nm.4236.

46. Walter, J., Britton, R.A., and Roos, S. (2011). Host-microbial symbiosis in the vertebrate gastrointestinal tract and the Lactobacillus reuteri paradigm. Proc Natl Acad Sci U S A 108 *Suppl 1*, 4645–4652. 10.1073/pnas.1000099107.

47. Spindler, M.P., Siu, S., Mogno, I., Li, Z., Yang, C., Mehandru, S., Britton, G.J., and Faith, J.J. (2022). Human gut microbiota stimulate defined innate immune responses that vary from phylum to strain. Cell Host Microbe 30, 1481–1498 e1485. 10.1016/j.chom.2022.08.009.

48. Vilela, J.A.P., Perrechil, F.A., Picone, C.S.F., Sato, A.C.K., and Cunha, R.L.D. (2015). Preparation, characterization and in vitro digestibility of gellan and chitosan-gellan microgels. Carbohydr Polym 117, 54–62. 10.1016/j.carbpol.2014.09.019.

49. Wastyk, H.C., Fragiadakis, G.K., Perelman, D., Dahan, D., Merrill, B.D., Yu, F.B., Topf, M., Gonzalez, C.G., Van Treuren, W., Han, S., et al. (2021). Gut-microbiota-targeted diets modulate human immune status. Cell 184, 4137–4153 e4114. 10.1016/j.cell.2021.06.019.

50. Derrien, M., Belzer, C., and de Vos, W.M. (2017). Akkermansia muciniphila and its role in regulating host functions. Microb Pathog 106, 171–181. 10.1016/j.micpath.2016.02.005.

51. Gutiérrez-Alzate, K., Beltrán-Cotta, L.A., Rekowsky, B.S.D., Cavalheiro, C.P., and da Costa, M.P. (2024). Micro- and Nanoencapsulation of Probiotics: Exploring Their Impact on Animal-Origin Foods. Acs Food Sci Technol 4, 2799–2812. 10.1021/acsfoodscitech.4c00776.

52. Blanco-Miguez, A., Beghini, F., Cumbo, F., McIver, L.J., Thompson, K.N., Zolfo, M., Manghi, P., Dubois, L., Huang, K.D., Thomas, A.M., et al. (2023). Extending and improving metagenomic taxonomic profiling with uncharacterized species using MetaPhlAn 4. Nat Biotechnol 41, 1633–1644. 10.1038/s41587-023-01688-w.

53. Beghini, F., McIver, L.J., Blanco-Miguez, A., Dubois, L., Asnicar, F., Maharjan, S., Mailyan, A., Manghi, P., Scholz, M., Thomas, A.M., et al. (2021). Integrating taxonomic, functional, and strain-level profiling of diverse microbial communities with bioBakery 3. Elife 10. 10.7554/eLife.65088.

54. McMurdie, P.J., and Holmes, S. (2013). phyloseq: an R package for reproducible interactive analysis and graphics of microbiome census data. PLoS One 8, e61217. 10.1371/journal.pone.0061217.

55. Excoffier, L., Smouse, P.E., and Quattro, J.M. (1992). Analysis of molecular variance inferred from metric distances among DNA haplotypes: application to human mitochondrial DNA restriction data. Genetics 131, 479–491. 10.1093/genetics/131.2.479.

56. Anderson, M.J. (2001). A new method for non-parametric multivariate analysis of variance. Austral Ecol 26, 32–46. DOI 10.1111/j.1442-9993.2001.01070.pp.x.

57. Bankhead, P., Loughrey, M.B., Fernandez, J.A., Dombrowski, Y., McArt, D.G., Dunne, P.D., McQuaid, S., Gray, R.T., Murray, L.J., Coleman, H.G., et al. (2017). QuPath: Open source software for digital pathology image analysis. Sci Rep 7, 16878. 10.1038/s41598-017-17204-5.

58. Galaxy, C. (2022). The Galaxy platform for accessible, reproducible and collaborative biomedical analyses: 2022 update. Nucleic Acids Res 50, W345–W351. 10.1093/nar/gkac247.

59. Subramanian, A., Tamayo, P., Mootha, V.K., Mukherjee, S., Ebert, B.L., Gillette, M.A., Paulovich, A., Pomeroy, S.L., Golub, T.R., Lander, E.S., and Mesirov, J.P. (2005). Gene set enrichment analysis: a knowledge-based approach for interpreting genome-wide expression profiles. Proc Natl Acad Sci U S A 102, 15545–15550. 10.1073/pnas.0506580102.

60. Mootha, V.K., Lindgren, C.M., Eriksson, K.F., Subramanian, A., Sihag, S., Lehar, J., Puigserver, P., Carlsson, E., Ridderstrale, M., Laurila, E., et al. (2003). PGC-1alpha-responsive genes involved in oxidative phosphorylation are coordinately downregulated in human diabetes. Nat Genet 34, 267–273. 10.1038/ng1180.

