## Supplementary figures and images for "Systemic Cysteine Elevation Sustains T-Cell Activation to Potentiate PD-1 Blockade"

### Graphical abstract

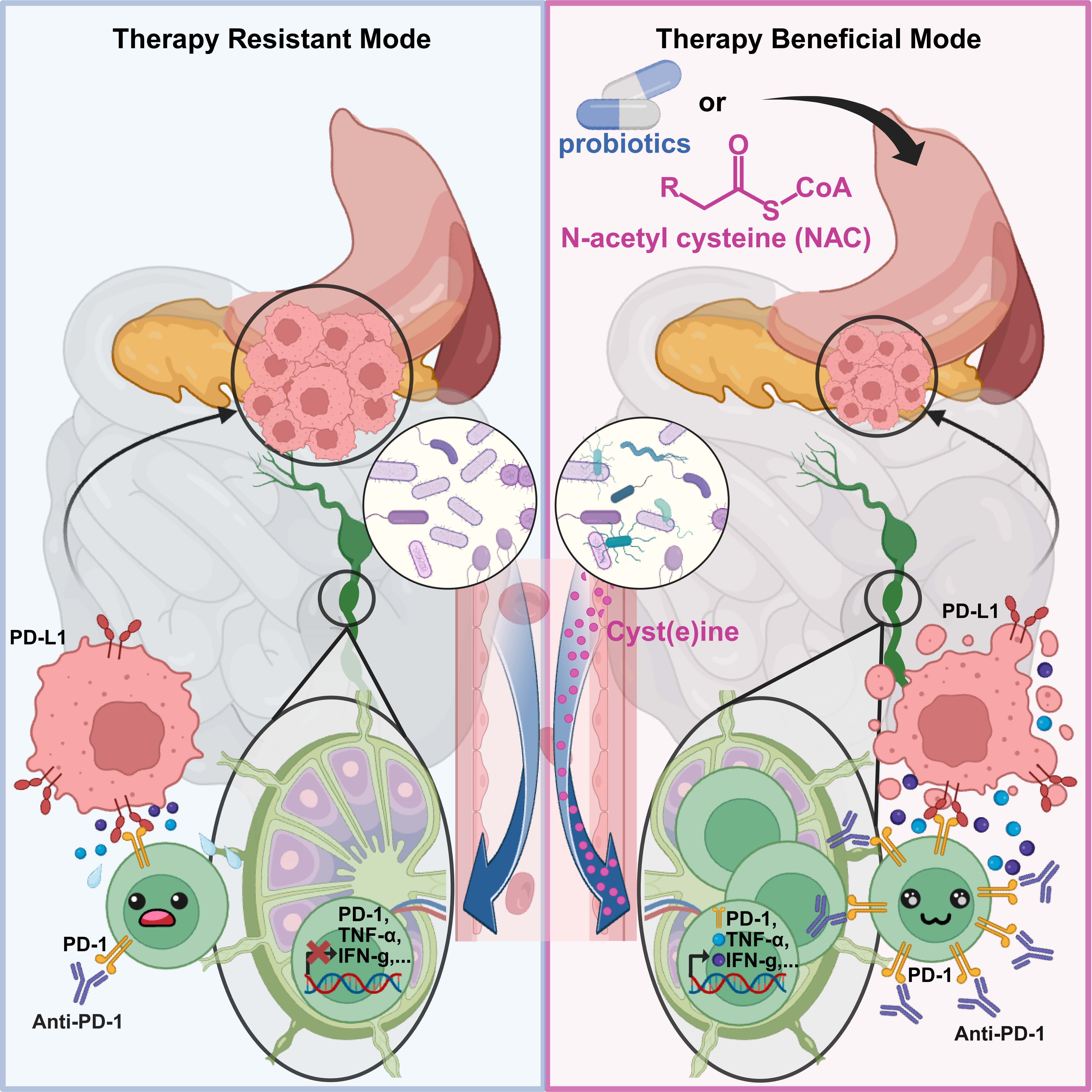

### Supplemental Figure 1

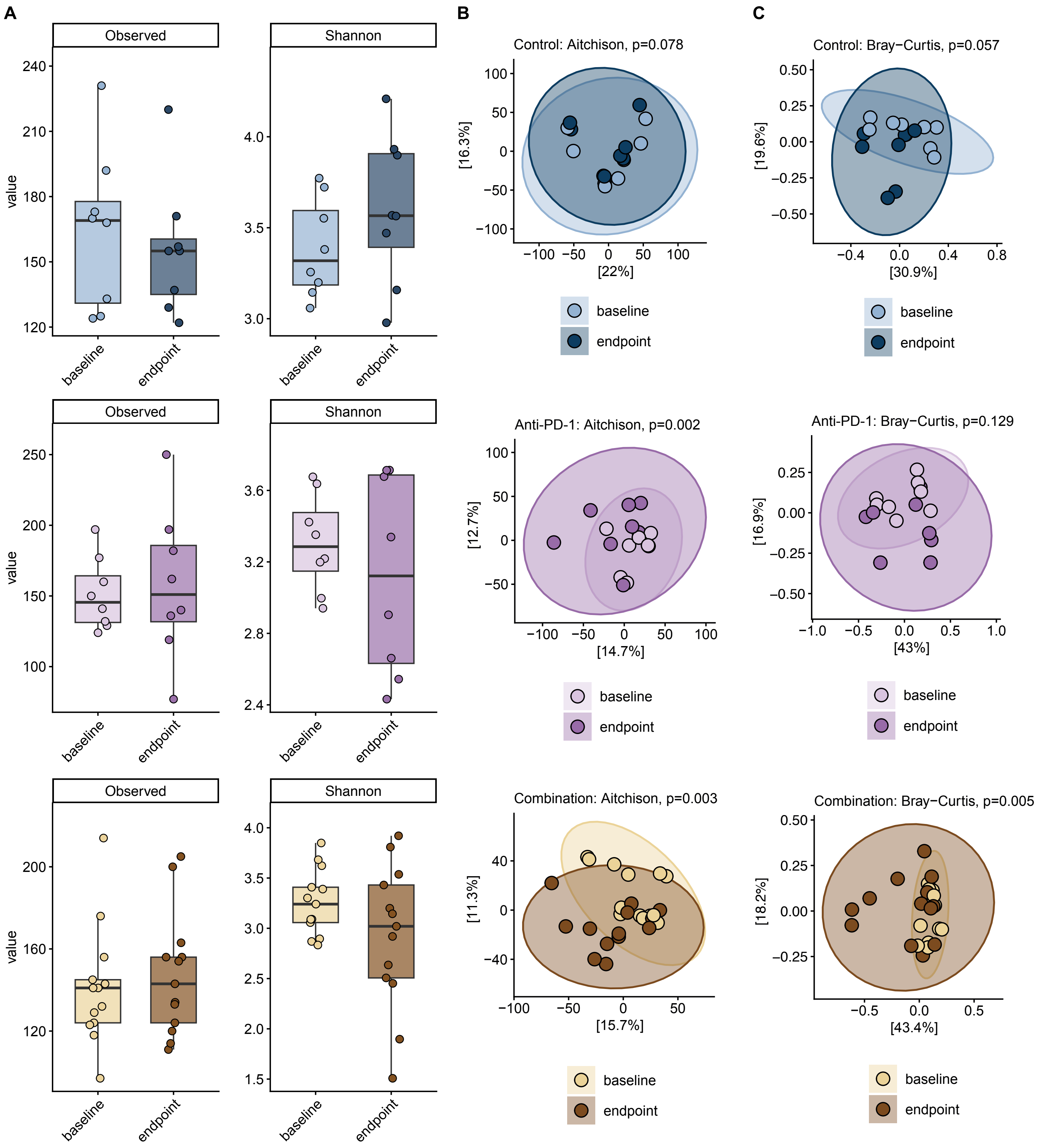

### Supplemental Figure 2

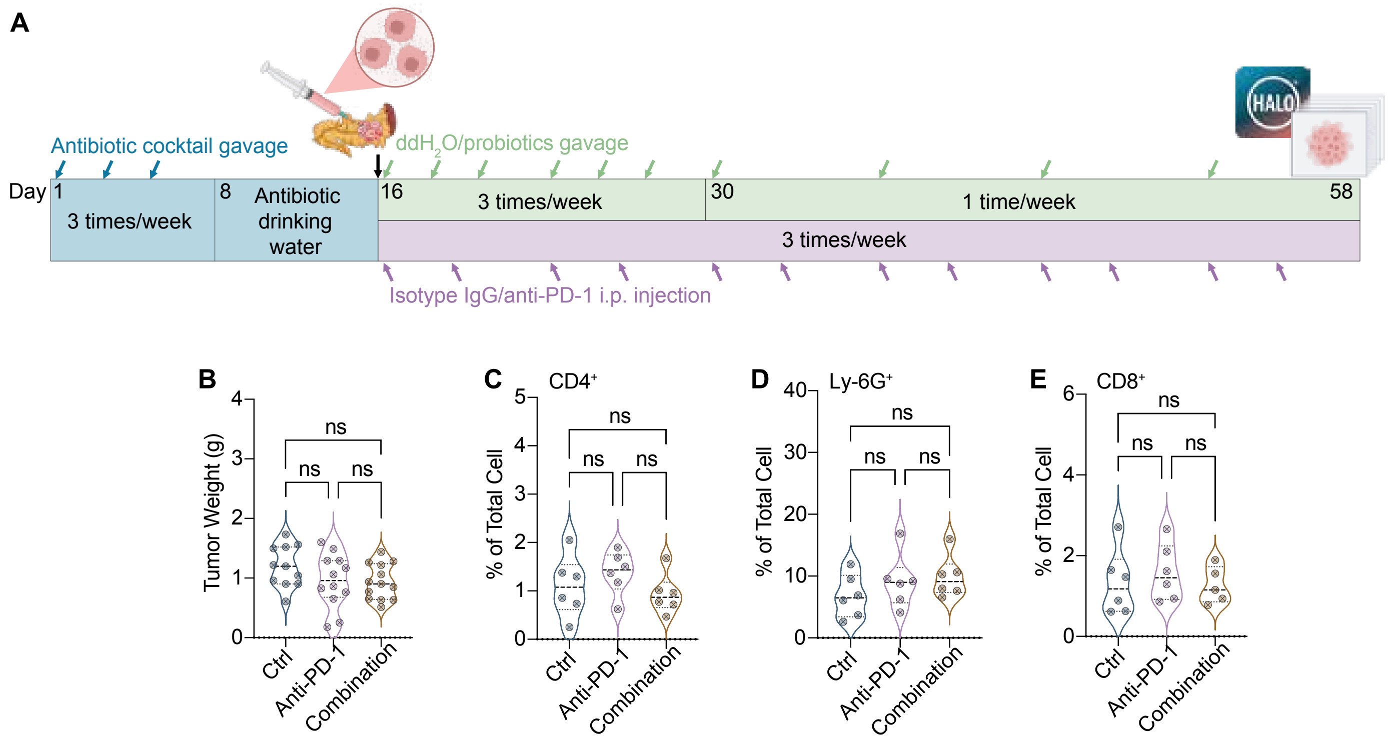

### Supplemental Figure 3

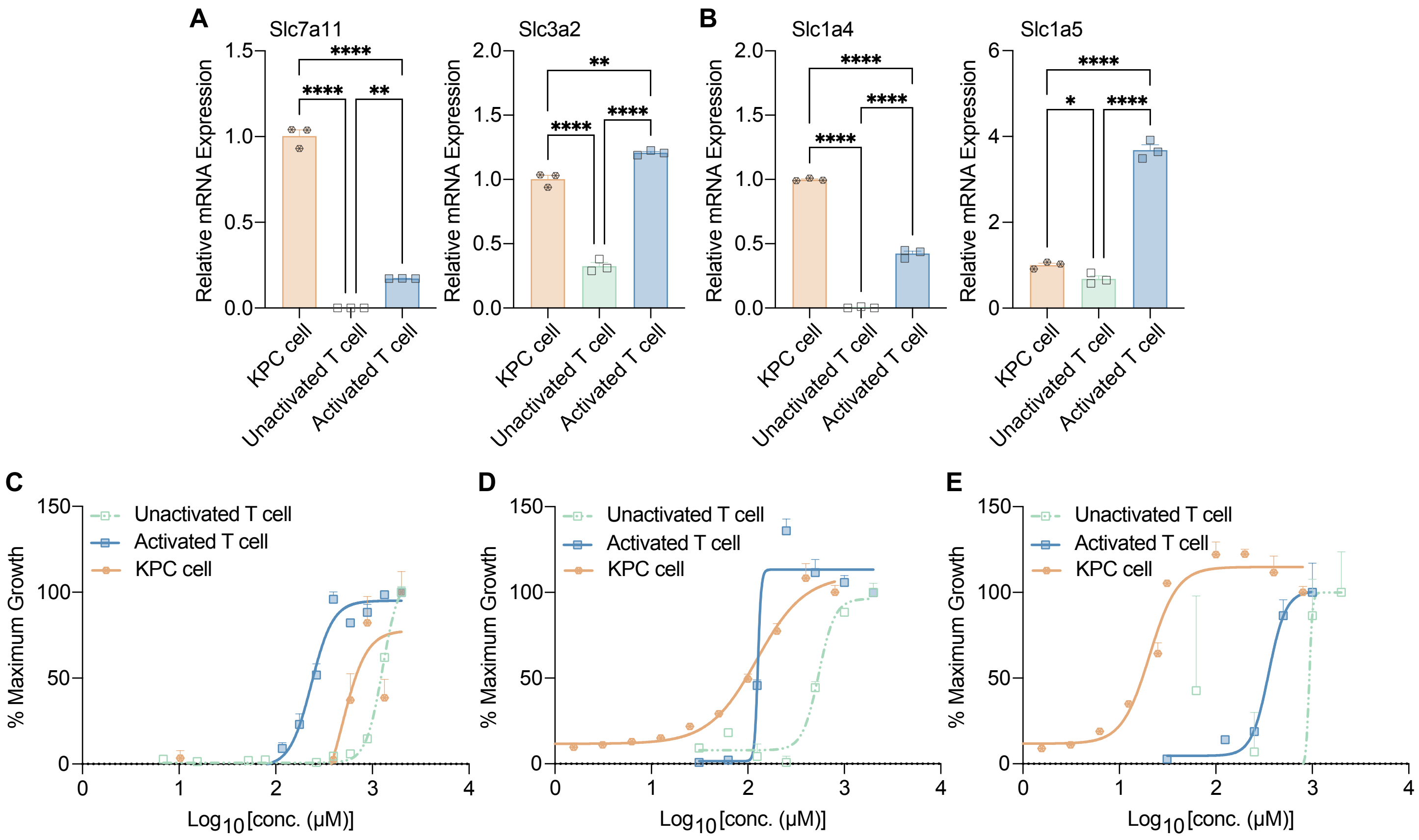

### Supplemental Figure 4

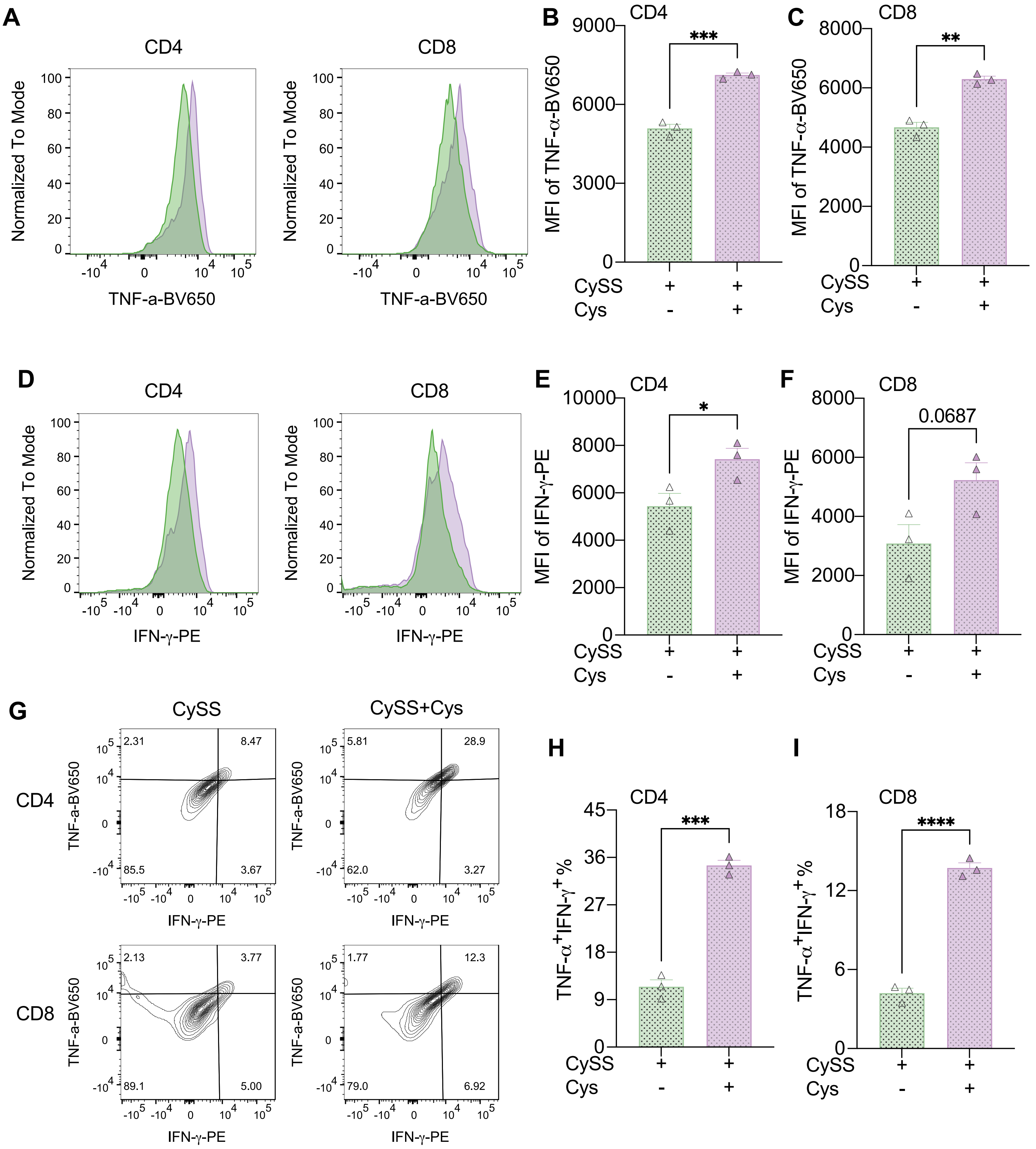

### Supplemental Figure 5

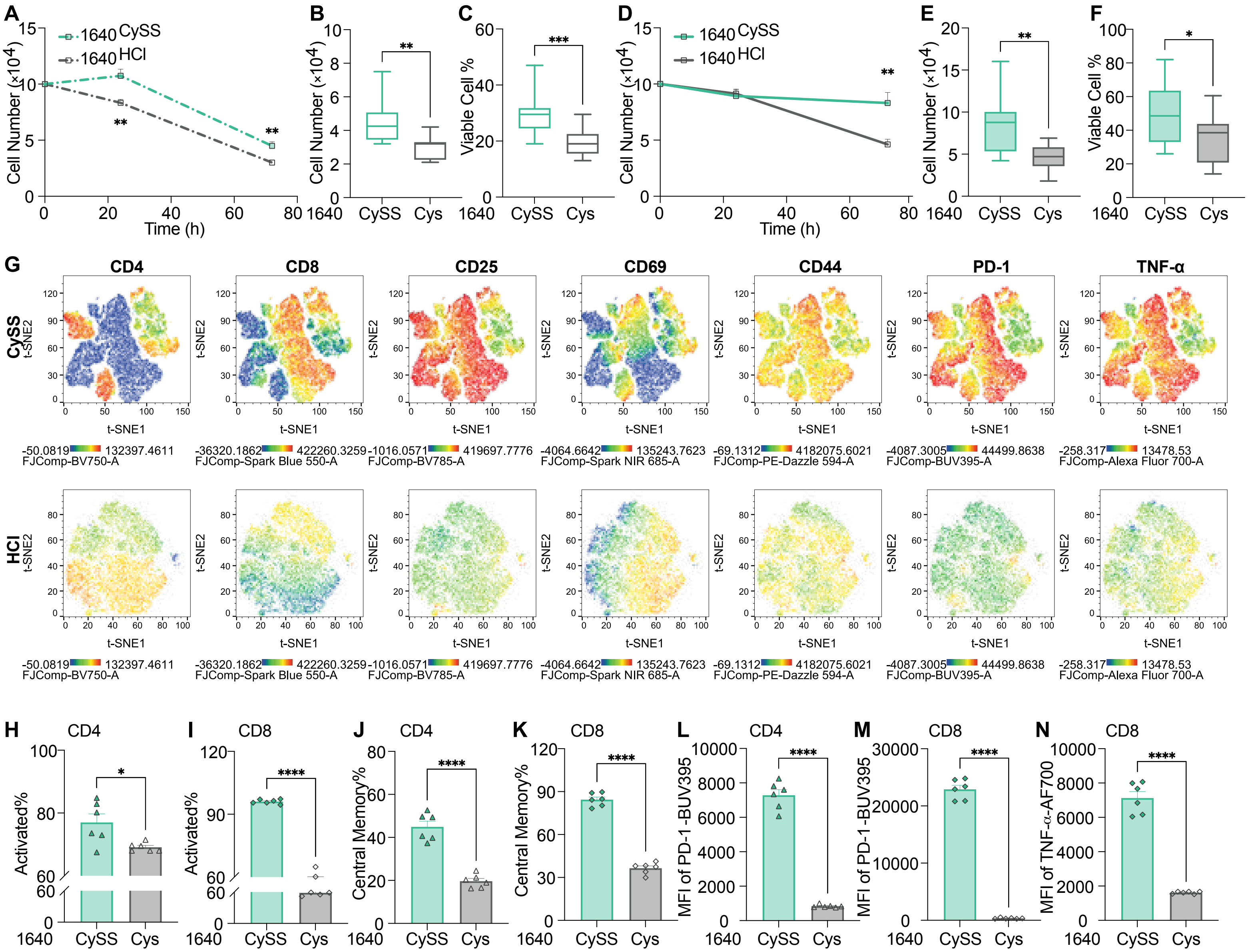

### Supplemental Figure 6

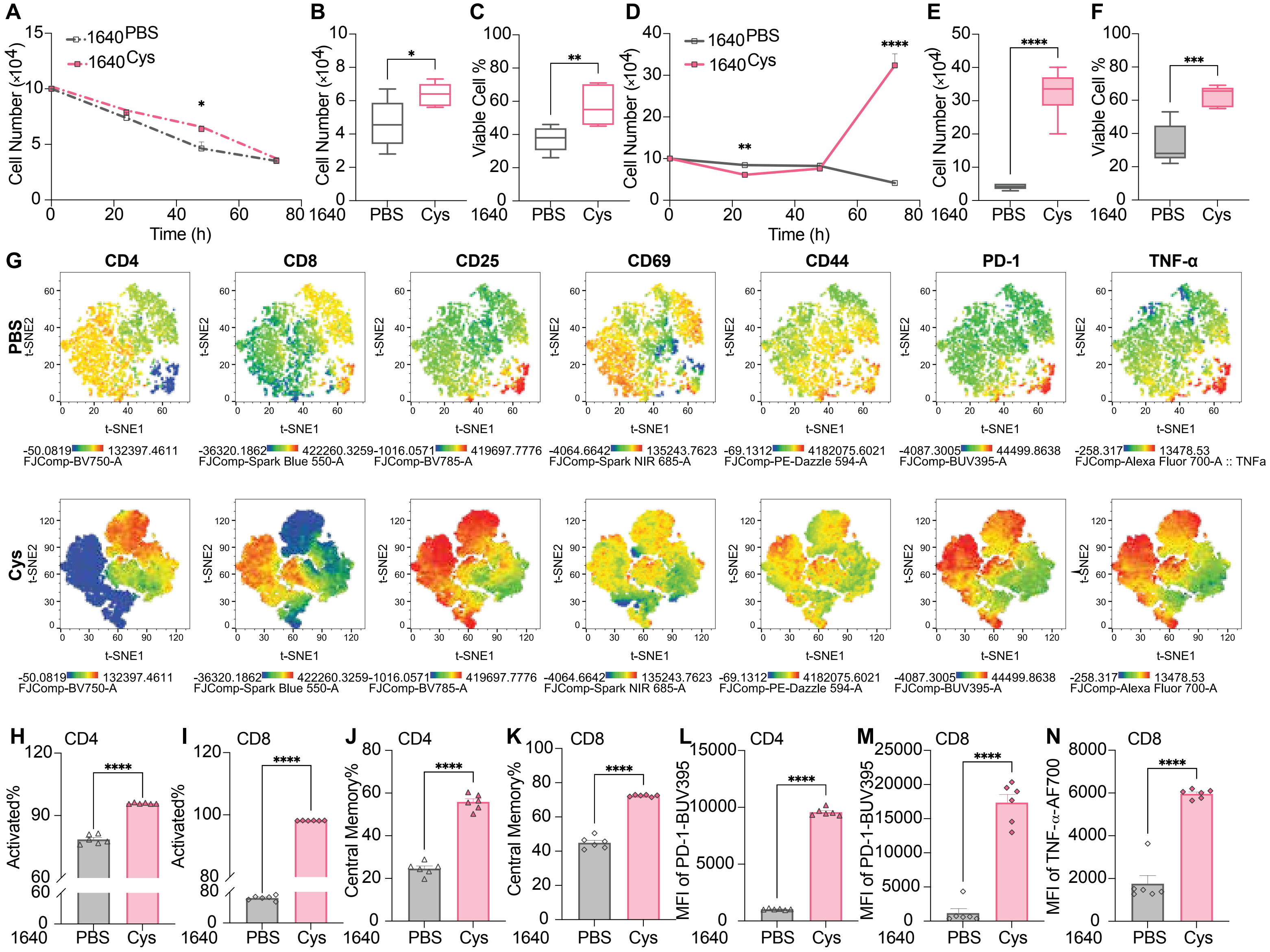

### Supplemental Figure 7

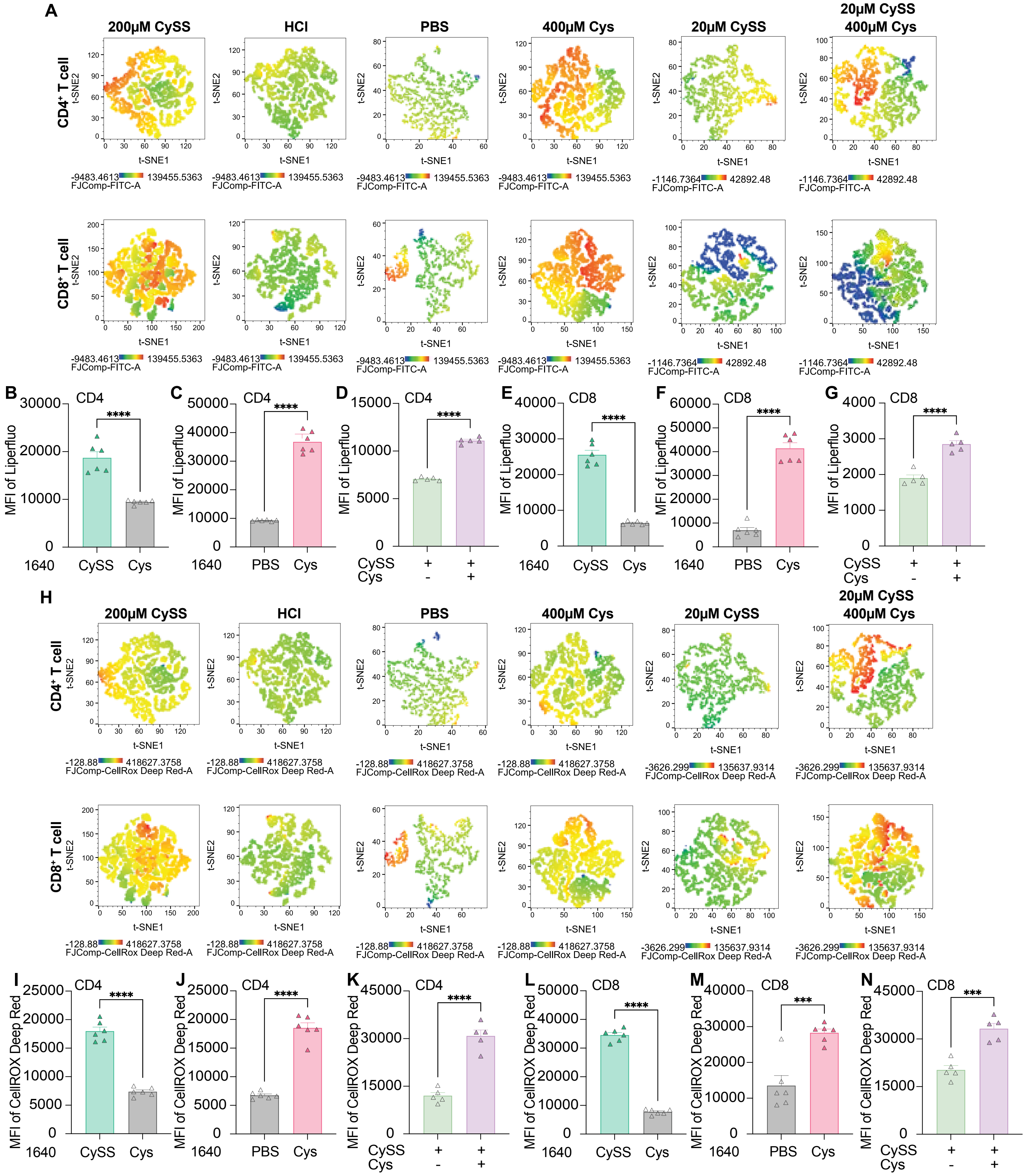
