## Supplemental Table 2-5 for "Systemic Cysteine Elevation Sustains T-Cell Activation to Potentiate PD-1 Blockade"

Table S2. Significant MetaCyc pathways identified by Wilcoxon testing ($p<0.05$), comparing the delta change in pathway CPM between the Anti-PD-1 and Combination groups. Related to Figure 3.

| PathwayID | p_value | mean_group1 | mean_group2 | fdr | n_group1 | n_group2 | comparison | Pathway |
|---|---|---|---|---|---|---|---|---|
| ASPASN-PWY | 0.02461055 | -15.897638 | 7.41216154 | 0.66279619 | 8 | 13 | αPD-1 vs Combination | superpathway of L-aspartate and L-asparagine biosynthesis |
| GLCMANNANAUT-PWY | 0.00993661 | -33.88675 | 6.25366923 | 0.63905843 | 8 | 13 | αPD-1 vs Combination | superpathway of N-acetylglucosamine, N-acetylmannosamine and N-acetylneuraminate degradation |
| PWY-1042 | 0.01989287 | 58.113 | -32.706692 | 0.63905843 | 8 | 13 | αPD-1 vs Combination | glycolysis IV |
| PWY-5667 | 0.01989287 | 69.6225375 | -11.060292 | 0.63905843 | 8 | 13 | αPD-1 vs Combination | CDP-diacylglycerol biosynthesis I |
| PWY-6293 | 0.01704155 | -2.6696163 | 1.64368462 | 0.63905843 | 8 | 13 | αPD-1 vs Combination | superpathway of L-cysteine biosynthesis (fungi) |
| PWY-6629 | 0.04455256 | -74.0156 | 27.3300615 | 0.74585163 | 8 | 13 | αPD-1 vs Combination | superpathway of L-tryptophan biosynthesis |
| PWY-724 | 0.04455256 | 45.24825 | 11.8834615 | 0.74585163 | 8 | 13 | αPD-1 vs Combination | superpathway of L-lysine, L-threonine and L-methionine biosynthesis II |
| PWY-7323 | 0.02578974 | 2.88907125 | -2.0757531 | 0.66279619 | 8 | 13 | αPD-1 vs Combination | superpathway of GDP-mannose-derived O-antigen building blocks biosynthesis |
| PWY-801 | 0.01704155 | -1.6212263 | 0.99745231 | 0.63905843 | 8 | 13 | αPD-1 vs Combination | homocysteine and cysteine interconversion |
| PWY0-1261 | 0.04565225 | -5.3548838 | 0.53123462 | 0.74585163 | 8 | 13 | αPD-1 vs Combination | anhydromuropeptides recycling I |
| PWY0-1297 | 0.04455256 | -28.143675 | -6.2643985 | 0.74585163 | 8 | 13 | αPD-1 vs Combination | superpathway of purine deoxyribonucleosides degradation |
| PWY0-1319 | 0.01989287 | 69.6225375 | -11.060292 | 0.63905843 | 8 | 13 | αPD-1 vs Combination | CDP-diacylglycerol biosynthesis II |
| RHAMCAT-PWY | 0.01989287 | 27.0692125 | -11.587922 | 0.63905843 | 8 | 13 | αPD-1 vs Combination | L-rhamnose degradation I |
| TRPSYN-PWY | 0.00774485 | -66.91385 | 21.9175 | 0.63905843 | 8 | 13 | αPD-1 vs Combination | L-tryptophan biosynthesis |

Table S3. Cyst(e)ine transporters in qPCR analysis. Related to Figure S3.

| Category | Sodium Dependence | System | Genes | Substrates | Detectable in T cells? | Detectable in KPC cells? |
|---|---|---|---|---|---|---|
| Acidic amino acid transporters | Sodium Dependent | $xAG^-$ | Slc1a1 | CySS, Cys | No | No |
| | Sodium Independent | $xc^-$ | Slc7a11 | CySS | Yes | Yes |
| Neutral amino acid transporters | Sodium Dependent | $y^+L$ | Slc3a2 | CySS | Yes | Yes |
| | | ASC | Slc1a4 | Cys | Yes | Yes |
| | | | Slc1a5 | Cys | Yes | Yes |
| | Sodium Independent | $b^{0,+}$ | Slc7a9 | Cys | No | No |
| | | ASC | Slc7a10 | Cys | No | Yes |

Table S4. Half-maximal growth concentration ($EC_{50}$) of cyst(e)ine in unactivated T cells, activated T cells, and KPC cells. Related to Figure S3.

| | $EC_{50}$ (μM) | | |
|---|---|---|---|
| | Cys (CySS based) | Cys (PBS based) | CySS (HCl based) |
| Unactivated T cell | 1276.0 | 541.8 | 929.3 |
| Activated T cell | 234.1 | 127.1 | 353.2 |
| KPC cell | 489.0 | 124.0 | 20.60 |

Table S5. Numbers and percentages of positively and negatively enriched pathways in each hierarchy following cysteine supplementation. Related to Figure 5.

| Positively Enriched | | | | Negatively Enriched | | | |
|---|---|---|---|---|---|---|---|
| Hierarchy | | Percentage (%) | Number | Hierarchy | | Percentage (%) | Number |
| Cell Cycle | | 28.80 | 53 | Cell Cycle | | 1.61 | 4 |
| DNA Replication and Repair | Chromatin Organization | 4.35 | 8 | DNA Replication and Repair | Chromatin Organization | 0.00 | 0 |
| | DNA Repair | 16.30 | 30 | | DNA Repair | 0.00 | 0 |
| | DNA Replication | 1.63 | 3 | | DNA Replication | 0.00 | 0 |
| Autophagy | | 0.00 | 0 | Autophagy | | 2.01 | 5 |
| Programmed Cell Death | | 1.09 | 2 | Programmed Cell Death | | 2.01 | 5 |
| Signal Transduction | | 4.35 | 8 | Signal Transduction | | 28.51 | 71 |
| Cell-Cell Communication | | 0.00 | 0 | Cell-Cell Communication | | 1.20 | 3 |
| Cellular Responses to Stimuli | | 3.26 | 6 | Cellular Responses to Stimuli | | 3.21 | 8 |
| Immune System | | 1.09 | 2 | Immune System | | 22.89 | 57 |
| Metabolism | | 11.96 | 22 | Metabolism | | 4.82 | 12 |
| Metabolism of RNA | | 5.43 | 10 | Metabolism of RNA | | 2.01 | 5 |
| Metabolism of Proteins | | 4.89 | 9 | Metabolism of Proteins | | 7.23 | 18 |
| Gene Expression (Transcription) | | 8.15 | 15 | Gene Expression (Transcription) | | 2.81 | 7 |
| Organelle Biogenesis and Maintenance | | 1.63 | 3 | Organelle Biogenesis and Maintenance | | 0.00 | 0 |
| Vesicle-Mediated Transport | | 1.63 | 3 | Vesicle-Mediated Transport | | 6.43 | 16 |
| Other | Hemostasis | 1.63 | 3 | Other | Hemostasis | 3.21 | 8 |
| | Muscle Contraction | 0.54 | 1 | | Muscle Contraction | 0.40 | 1 |
| | Neuronal System | 2.72 | 5 | | Neuronal System | 0.80 | 2 |
| | Reproduction | 0.54 | 1 | | Developmental Biology | 4.82 | 12 |
| | | | | | Extracellular Matrix Organization | 1.20 | 3 |
| | | | | | Transport of Small Molecules | 4.82 | 12 |
